# Prophylactic host behaviour discourages pathogen exploitation

**DOI:** 10.1101/2019.12.18.881383

**Authors:** Evan Mitchell, Geoff Wild

## Abstract

Much work has considered the evolution of pathogens, but little is known about how they respond to changes in host behaviour. We build a model where hosts are able to choose to engage in prophylactic measures that reduce the likelihood of disease transmission. This choice is mediated by costs and benefits associated with prophylaxis, but the fraction of hosts engaged in prophylaxis is also affected by population dynamics. We identify a critical cost threshold above which hosts do not engage in prophylaxis. Below the threshold, prophylactic host behaviour does occur and pathogen virulence, measured by the extent to which it exploits its host, is reduced by the action of selection relative to the level that would otherwise be predicted in the absence of prophylaxis. Our work emphasizes the significance of the dual nature of the trade-off faced by the pathogen between balancing transmission and recovery, and creating new infections in hosts engaging or not engaging in prophylaxis.

## 1. Introduction

Mathematical study of infectious diseases has a long history that predates even the well-known contributions of Ross [1] and Kermack and McKendrick [2]. Mathematical study of the *evolution* of infectious disease-causing agents, however, is relatively recent. One avenue of inquiry has explored the ways in which natural selection shapes the virulent effects disease-causing agents (pathogens) inflict on their hosts. Models have provided insight into the ways in which various factors—such as parasite reproductive rates [3], host density [4], relatedness among co-infecting pathogen strains [5], and multiple types of hosts [6]—modulate the expressed level of virulence. A key prediction emerging from this body of work is that, in many cases, selection acts to maximize the basic reproduction number [7, 8], i.e., the expected number of new infections created by a single infective host [9]. In particular, adaptive levels of pathogen virulence balance a trade-off between the average duration of infection and the rate of disease transmission. As disease transmission rates slow, then, standard theory predicts selection will respond by shaping virulence in a way that decreases the duration of infection.

Mathematics has also contributed to our understanding of the ways in which host traits impact the evolution of pathogen virulence. Here, models have explored the effects of coevolution with innate host defences [8, 10], use of antibiotics [11], vaccines and vaccination behaviour [12, 13], and other social factors related to hosts themselves [14]. It is this last item—namely, the effect of host social behaviour on the evolution of pathogen virulence—on which we focus our attention in this paper. Schaller [15] discusses the idea of a “behavioural immune system” that complements the standard physiological immune system in humans. This behavioural immune system is comprised of various prophylactic measures, such as social distancing (e.g., avoiding handshakes when greeting) or improved personal hygiene (e.g., hand washing), that individuals may adopt to reduce the likelihood of infection [15].

Unfortunately, little is known about how hosts’ behavioural immune system impacts the evolution of pathogens. What work has been done in this area predicts that prophylactic behaviour exhibited by hosts can select for higher pathogen virulence [16]. This work, however, considers short-term evolutionary outcomes only, and does not completely link host behavioural changes with host demographics and disease dynamics. In order to fully assess the risks posed by pathogen evolution, then, more detailed models are required. We devise a model that tightly couples changes in the host’s behvioural immune system, host demographics, and disease dynamics, in a way that allows us to make predictions about the long-term evolution of pathogen virulence. In contrast to previous work, we find prophylactic behaviour uniformly reduces the pathogen’s evolutionarily stable level of virulence below the level expected in the absence of such behaviour. Our result is also surprising in light of the standard theory that associates increased virulence to decreases in the transmissibility of a disease [8]. We argue that the driving force behind our result is the gradual decoupling of the evolutionary and behavioural dynamics, mediated by the cost of prophylactic measures.

## 2. Model

### (a) Disease Dynamics

We begin with a version of an endemic SIR model of infectious-disease dynamics [17] modified in a way that separates a host population of total size *N* into two groups. Individuals in group *i* = 0 are those who do not take prophylactic measures that limit (but do not completely prevent) disease transmission, whereas those in group *i* = 1 do take such measures. Individuals in each group are further subdivided according to their disease status. Let *S*_*i*_, *I*_*i*_, and *R*_*i*_ denote the number of individuals in group *i* who are susceptible to infection, infective, and recovered from infection, respectively.

Each individual encounters another at a fixed rate. Given that an encounter is between a susceptible and an infective, the likelihood of disease transmission depends on the groups to which individuals belong. If *β*_*ij*_ denotes the product of the probability of disease transmission from a *j*-infective to an *i*-suceptible and the per-capita encounter rate, then *S*_*i*_*β*_*ij*_*I*_*j*_*/N* gives us the total rate at which new infections are created in group *i*. Infective individuals recover at a fixed per-capita rate, *γ*, independent of their group. As a result of recovery, individuals are imbued with life-long immunity to future infection. Individuals can also switch groups. For now we use *τ*_*ij*_, *ϕ*_*ij*_ and *η*_*ij*_ to represent the per-capita rates at which susceptible, infective, and recovered individuals, repsectively, switch from group *j* to *i*. We expand on the details surrounding group switching later. We can summarize the description above using a system of differential equations. Scaling time so that one time unit is equivalent to the average lifetime of an individual in the population, and matching birth and death rates, we get

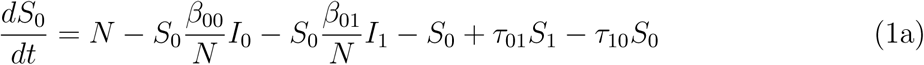

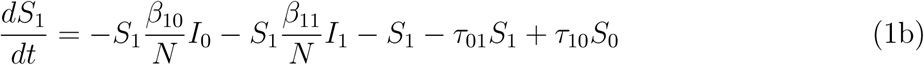

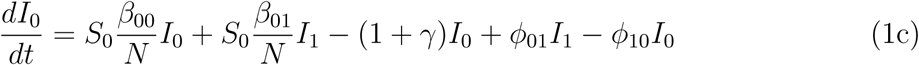

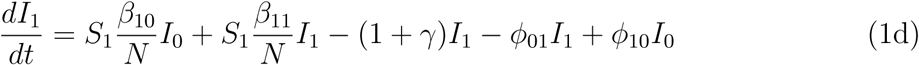

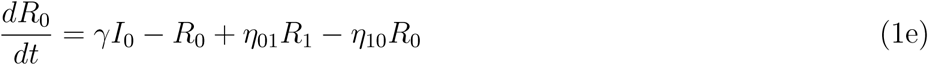

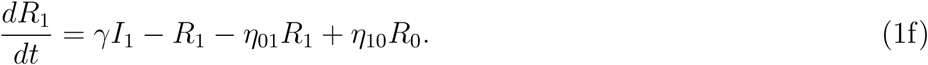

Note that the differential equations in (1) sum to zero, and so total population size *N* is constant. This, along with the fact that group membership among recovered individuals is of no consequence, allows us to omit (1e) and (1f). We now use *u*_*i*_ = *S*_*i*_*/N* and *v*_*i*_ = *I*_*i*_*/N* to denote the fraction of susceptible and infective individuals, respectively, in group *i*. Similarly, we use *u* = *u*_0_ + *u*_1_ and *v* = *v*_0_ + *v*_1_ to denote the total fraction of susceptible and infective individuals, respectively. The dynamics of *u* and *v* can then be modelled by the following set of differential equations:

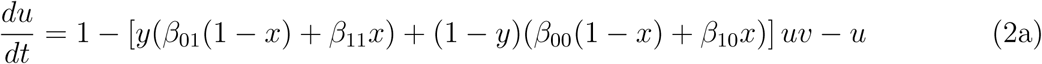

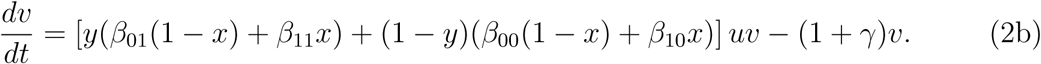

We also track the fraction of susceptible and infected individuals taking protective measures using *x* = *u*_1_*/u* and *y* = *v*_1_*/v*, respectively. Given this definition of *x*, we can derive the following differential equation to describe how the proportion of susceptible individuals engaging in prophylaxis changes due to various factors:

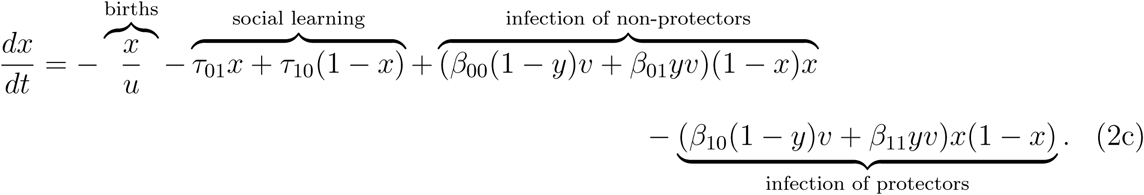

The first term represents the fact that births increase the pool of non-protectors (i.e., individuals not engaging in prophylactic measures), which in turn reduces the relative proportion of susceptible protectors (i.e., individuals engaging in prophylactic measures). The *τ*_*ij*_ terms capture the effects of individuals switching between the two classes of susceptibles as a result of the costs and benefits of prophylaxis. The final two terms correspond to infection. In one case, infection reduces the pool of non-protectors and subsequently increases the relative proportion of protectors. Conversely, those engaging in prophylaxis can also become infected, reducing the proportion of protectors. Similarly, we can define the differential equation for *y* as:

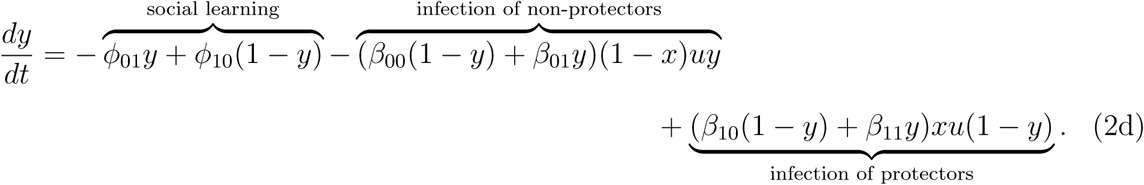

The first term represents the effects of individuals switching between infective classes due to the costs and benefits of prophylaxis. The latter two terms correspond to infection in the same vein as in equation (2c).

Equations (2c) and (2d) capture dynamic features of the protector population that previous work has ignored. Modelling undertaken by Pharaon and Bauch [16] accounts for social learning but neglects changes due to demographics (births) and infection. Our equations, therefore, combine multiple processes to establish a more realistic model of changing host behaviour.

We now return to the issue of group switching, and the details surrounding *τ*_*ij*_ and *ϕ*_*ij*_. Individuals do not always adhere to beneficial measures such as taking medications [18], exercise regimes [19], or dietary restrictions [20], so we include group switching in our model to account for these types of effects. Following [16], we use the replicator dynamic [21] to model the total rate at which individuals move from one group to another. We assume that the decision to switch groups is driven by utility, as discussed in [22]. Specifically, an individual’s utility will be determined by their risk of infection less the cost of taking prophylactic measures. For an *i*-susceptible, the risk of infection it faces will be quantified by the force of infection,

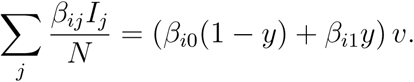

Consequently, the utility of a susceptible individual adopting prophylactic measures is

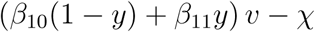

where *χ* is the cost to engaging in prophylaxis. Infective individuals have no risk of infection, so they only pay the cost of the prophylactic measure should they choose to engage in it. Thus, *-χ* represents the utility of an infective individual engaging in prophylaxis. A susceptible (resp. infective) individual will then choose to switch into a given group when the utility of an individual in that group exceeds the utility of the average susceptible (resp. infective) in the population. Ultimately, the replicator-dynamics model of switching gives us

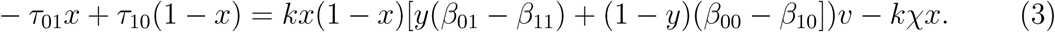

and

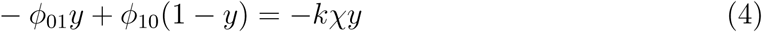

where *k* is a constant that reflects the rate of social learning. We note that our model of switching makes the simplification (made elsewhere [16]) of assuming individuals have up-to-date information about the global state of the population. Our final model of disease dynamics can now be stated as,

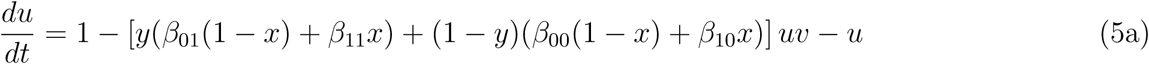

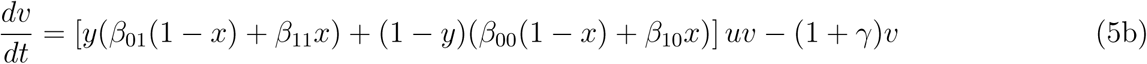

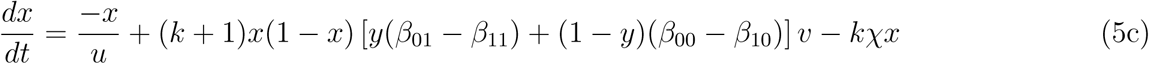

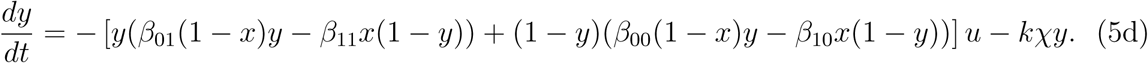

Note that, if no individual in the population takes protective measures, then *x* = *y* = 0 and we recover the standard endemic SIR model (Appendix A). When some fraction of the population takes protective measures, however, standard predictions of the SIR model may or may not hold. The linear stability analysis presented in Appendix A shows that the well-known endemic equilibrium remains stable as long as

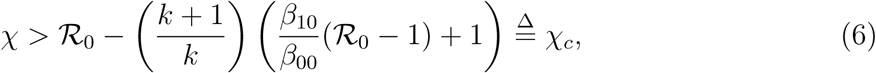

where 𝓡_0_ = *β*_00_*/*(1 + *γ*) is the basic reproductive number [9] of the system in the absence of protective measures. Below this threshold, the cost to taking protective measures is low enough that the system moves towards a new endemic equilibrium 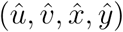 at which some non-zero fractions of susceptible and infective individuals adopt prophylactic measures. It is this new endemic equilibrium that will frame the evolutionary model we pursue in the next section.

### (b) Evolutionary Dynamics

We consider the evolution of the level of pathogen exploitation of its host, denoted *ξ >* 0. Exploitation affects disease transmission, with a greater *ξ* value corresponding to a greater *β*_*ij*_. To reflect this, we now write *β*_*ij*_(*ξ*) where

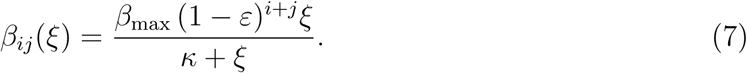

In words, we are treating *β*_*ij*_ as an increasing function of *ξ* that saturates at a value of *β*_max_(1 *- ε*)^*i*+*j*^, where *ε* is the probability that the prophylactic measures prevent disease transmission. We assume that prophylactic measures taken by individuals fail independently, so (1 *- ε*)^2^ gives the probability that the disease is transmitted between two individuals engaging in prophylaxis. The rate at which transmission saturates is controlled by *κ >* 0, with larger values of this constant corresponding to a reduced rate of saturation.

Exploitation also affects recovery. To reflect this assumption, we write *γ*(*ξ*), where *γ*(*ξ*) = *cξ* for a constant *c* with units of inverse time (the exploitation level *ξ* is dimensionless). Without loss of generality, we take *c* = 1. Here, increased exploitation acts to reduce the expected duration 1*/*(1 + *γ*(*ξ*)) of an infection. This penalty of larger *ξ*, then, trades off against the transmissibility benefits described above. Previous authors have either assumed (or shown) that such trade-offs exist, though they are often mediated by disease-related mortality [7, 23, 24] or viral load [25]. Here, we follow Úbeda and Jansen [26] and Alizon [27] by assuming the trade-off faced by the pathogen involves recovery, e.g., through increased viral load making it more likely that the pathogen is detected by the host’s immune system.

We use an adaptive dynamics approach to model the evolution of pathogen exploitation under the primary influence of natural selection [8, 28, 26]. We introduce a rare mutant pathogen with exploitation trait *ξ*_*m*_ into a resident pathogen population with exploitation trait *ξ*. It is assumed that the resident system has reached equilibrium prior to introducing the mutant, that there is no co-infection, and that the prophylactic measures are equally effective at preventing transmission of both strains.

Let *v*_*m*_ denote the fraction of individuals in the population infected with the mutant strain. While the mutant is rare, its dynamics are well approximated by

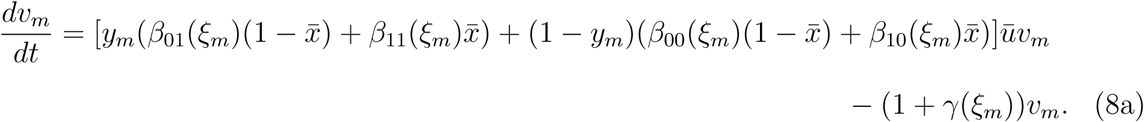

As we do with the resident population, we can also track the proportion of individuals infected with the mutant strain engaging in prophylaxis. Denoting this proportion by *y*_*m*_, we can describe its dynamics by

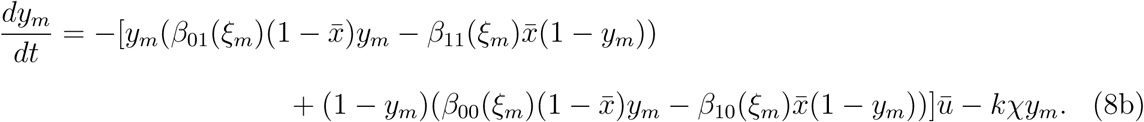

If the mutant strain becomes common, the approximate dynamics in (8) break down and we say that the mutant has successfully invaded the resident population. Provided the system is sufficiently close to an evolutionarily steady state, a mutant who successfully invades will become the new resident [29].

Since *v*_*m*_ does not appear in (8b), we can first solve for the equilibrium value 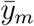 of *y*_*m*_, substitute that value into (8a), and study (8a) alone. If the right-hand side of (8a) is positive (resp. negative), the mutant invades (resp. is eliminated) because it is favoured (resp. disfavoured) by natural selection. The sign of the right-hand side of (8a) is the same as the sign of the difference between

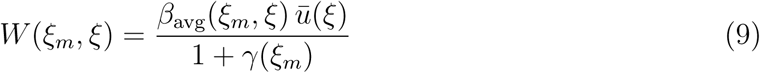

and unity, where

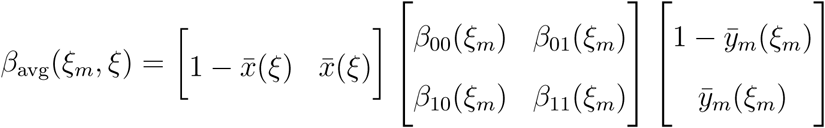

represents an average transmission rate taking into account the different groups of susceptible and infective individuals. *W* then has a clear biological interpretation, made in previous work [8], in terms of the basic reproductive number of the mutant strain. In particular, an infection with the mutant strain lasts an average of 1*/*(1 + *γ*) time units and an average of *β*_avg_*ū* new infections are created during this time. If this quantity is larger than one (resp. smaller than one), then the mutant population will grow (resp. shrink). Writing *W* as we have done in equation (9) also highlights the two trade-offs faced by the pathogen. One is the transmission-recovery trade-off described above, captured through the *β*_avg_ and *γ* terms. By unpacking the *β*_avg_ term, though, we can also see a second trade-off between the type of infection created. Through the 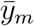 terms, the pathogen is able to influence whether a new infection occurs in a protector or a non-protector. We can, therefore, use *W* as a payoff function in the adaptive dynamics analysis.

Following the discussion above, the direction of evolution of *ξ* that is favoured by natural selection is given by the sign of 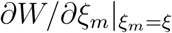. Consequently, the selective process is at equilibrium whenever 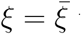 where 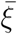 satisfies

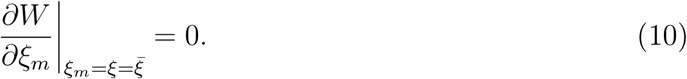

An equilibrium value 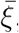, i.e., one that satisfies condition (10), may or may not be stable. If the equilibrium value 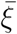 attracts nearby resident populations, then

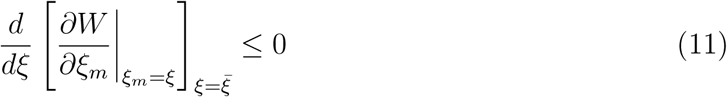

and we say that 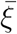 is convergence stable [30]. If the equilibrium 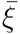 resists invasion from nearby mutants, then

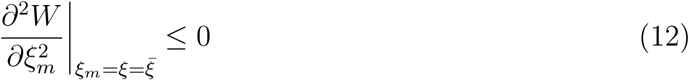

and we say that 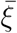 is evolutionarily stable (sensu [31]). A sufficient condition for either type of stability is obtained by replacing the weak inequality with a strict one. It is the strict sufficient versions that we use here. When 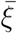 is both convergence stable and evolutionarily stable, we say it is a continuously stable strategy (CSS) [32].

## 3. Results

### (a) Simple Cases

There are two special cases that can be analyzed with relative ease. The first special case assumes the cost of prophylaxis exceeds the threshold *χ*_*c*_. In this case, no one in a population supporting the resident endemic disease is adopting prophylactic measures and so 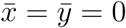. Given the absence of protectors, we find two equilibrium values of *y*_*m*_: a stable point 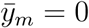 and an unstable point 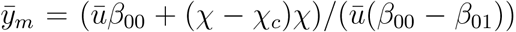. This, in turn, reduces the payoff function to

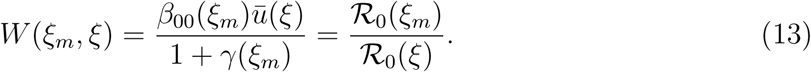

The mutant strain is then able to invade (resp. is eliminated) if *W* (*ξ*_*m*_, *ξ*) exceeds (resp. is less than) unity; equivalently, if 𝓡_0_(*ξ*_*m*_) exceeds or is less than 𝓡_0_(*ξ*). More importantly, the CSS level of exploitation, 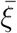, will maximize 𝓡_0_(*ξ*) and so 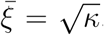. This will serve as a benchmark against which more general results will be compared.

The second special case assumes that prophylactic measures are cost-free, i.e., *χ* = 0. Under this assumption, the distribution of the mutant strain, captured by 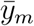, does not depend on the mutant exploitation level and so the trade-off in the type of infection created is no longer applicable. In this case, the payoff function simplifies substantially and analysis shows that 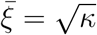 is the CSS. This is the same result as the benchmark established in our first special case, despite the presence of individuals engaging in prophylaxis. Absence of cost, it seems, decouples the pathogen’s evolution from the evolution of host behaviour due to the pathogen losing access to the trade-off in the type of infection created.

### (b) Evolution Near the Critical Cost

In general, the model cannot be explored analytically. However, if we are near the critical cost *χ*_*c*_ we can derive quasi-analytic results. When the cost *χ* is slightly below its critical threshold *χ*_*c*_, we can approximate the CSS exploitation level as 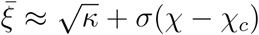 where *σ* is a constant such that

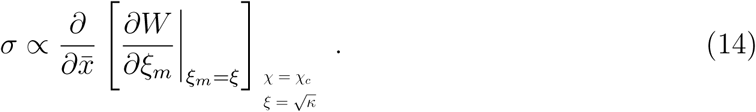

If *σ* is positive (resp. negative), then 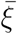 is above (resp. below) the benchmark value of 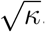. As equation (14) shows, whether we are above or below this benchmark depends on how small changes in cost lead to small changes in the proportion of susceptible protectors which, in turn, lead to changes in the selection gradient acting on exploitation.

We can show with a quasi-analytic approach that equation (14) is always negative. Our evidence relies on first choosing feasible values of our parameters. In particular, we need 𝓡_0_ *>* 1. If 𝓡_0_ *>* 1, then we need also to choose the probability 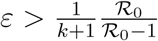, thus ensuring that *χ*_*c*_ *>* 0. To ensure that *ε <* 1, we then need to choose 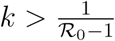.

Using feasible parameters and working to zeroth order in *χ − χ*_*c*_, we use the computer algebra software (CAS) Maple (version 2019.1) to investigate the sign of *σ* as described in equation (14). We find that the requirement that 𝓡_0_ *>* 1 necessarily restricts our choices of maximal transmissibility, *β*_max_, to values that lie above the curve traced out by 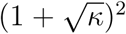 (figure 1, panel A). The CAS shows that nullclines of (14) never exceed the 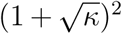 curve for the wide range of feasible parameters we investigated (figure 1, panel B). Thus, the sign of *σ* does not change provided the 𝓡_0_ *>* 1 restriction is met. Moreover, test points show that the sign of *σ* itself is negative when feasible model parameters are chosen. Based on CAS investigations described in Appendix C, then, we conclude that, just below the critical cost, selection acts to reduce the CSS level of host exploitation exhibited by the pathogen.

**Figure 1:**
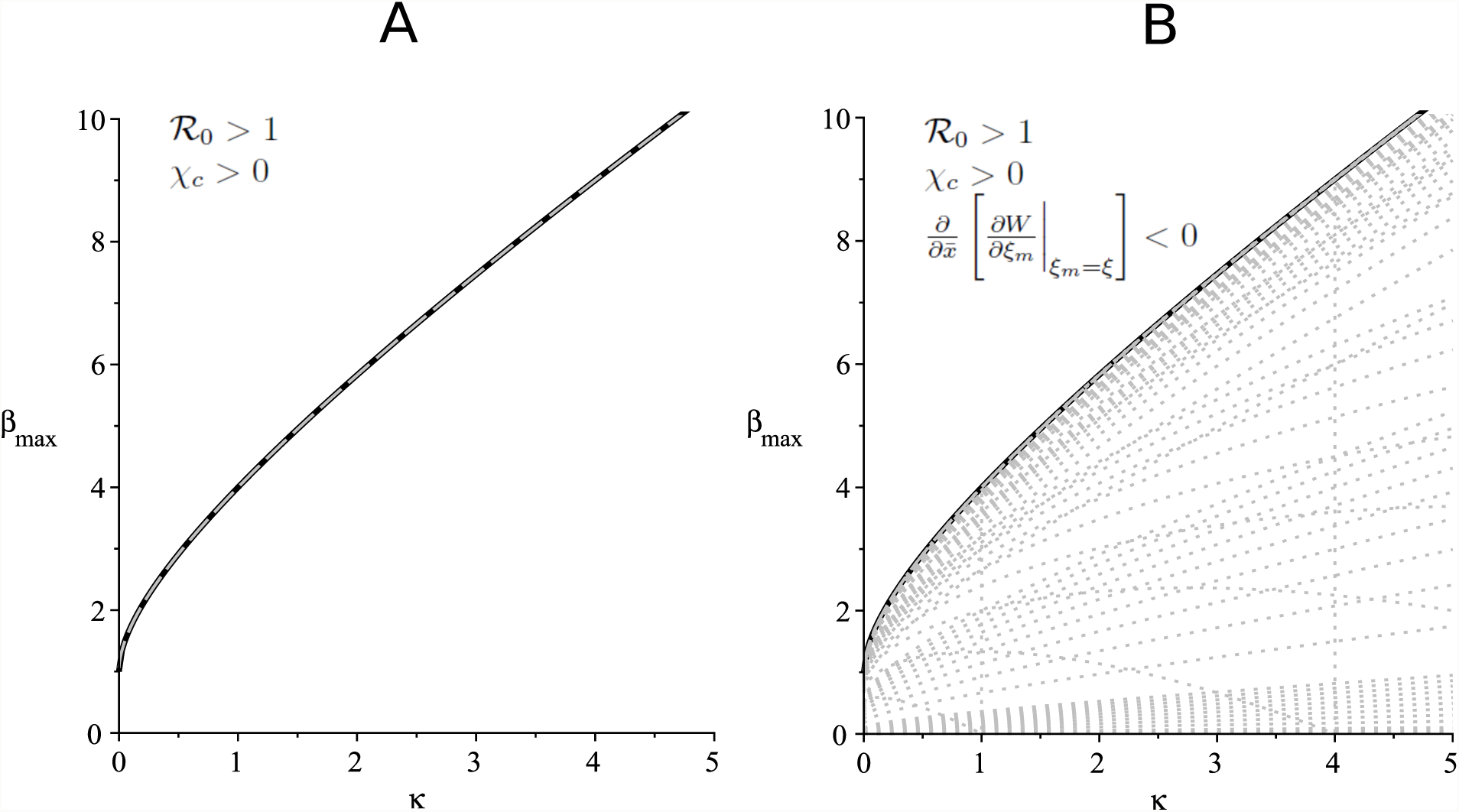
Panel A shows the curve *χ*_*c*_ = 0 (dashed grey) overlaying the curve 𝓡_0_ = 1 (solid black), and the region of parameter space where *χ*_*c*_ *>* 0 and 𝓡_0_ *>* 1. Dotted grey curves represent the roots of the right-hand side of equation (14) and panel B shows that these all occur on or below the black and dashed grey curves (note that vertical dotted grey lines are an artifact of jump discontinuities). Choosing parameter values in the region above the curves in panel A results in the right-hand side of equation (14) being negative, as noted in panel B. The implication is that the pathogen exploitation level (virulence) will decrease relative to the benchmark level of 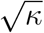 close to the cost threshold *χ*_*c*_.

### (c) Evolution for Arbitrary Cost

Our results can be extended numerically for costs that are possibly much smaller than the critical value, *χ*_*c*_, using a Matlab (version R2019a) procedure detailed in Appendix D. We build the procedure around the observation that locally asymptotically stable equilibrium solutions to 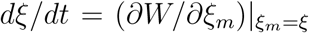 are also convergence-stable evolutionary equilibria as defined by conditions (10) and (11), respectively. As a result, numerical iteration of this differential equation can be used to find candidate CSS strategies. The evolutionary stability of candidate CSS strategies can be confirmed with a centred finite-difference approximation of (12). Since the error is on the order of the square of the distance between *ξ* values used in the approximation, we consider any value within this error to satisfy the ESS condition.

The results of our numerical procedure confirm that the benchmark CSS level of 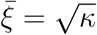 is obtained when *χ* = 0 and *χ* = *χ*_*c*_. Second, numerical results confirm the reduction in the CSS level of pathogen exploitation for *χ* slightly smaller than *χ*_*c*_. Third, and most important, numerical results indicate that the CSS exploitation rate 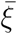 changes in a simple way as cost is reduced from its critical value to its natural lower limit at zero (figure 2). In particular, as cost is reduced 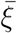 decreases monotonically from the benchmark value until it reaches a minimum. Once at the minimum, the direction of selection changes and 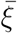 increases monotonically, ultimately returning to the benchmark when cost disappears.

**Figure 2:**
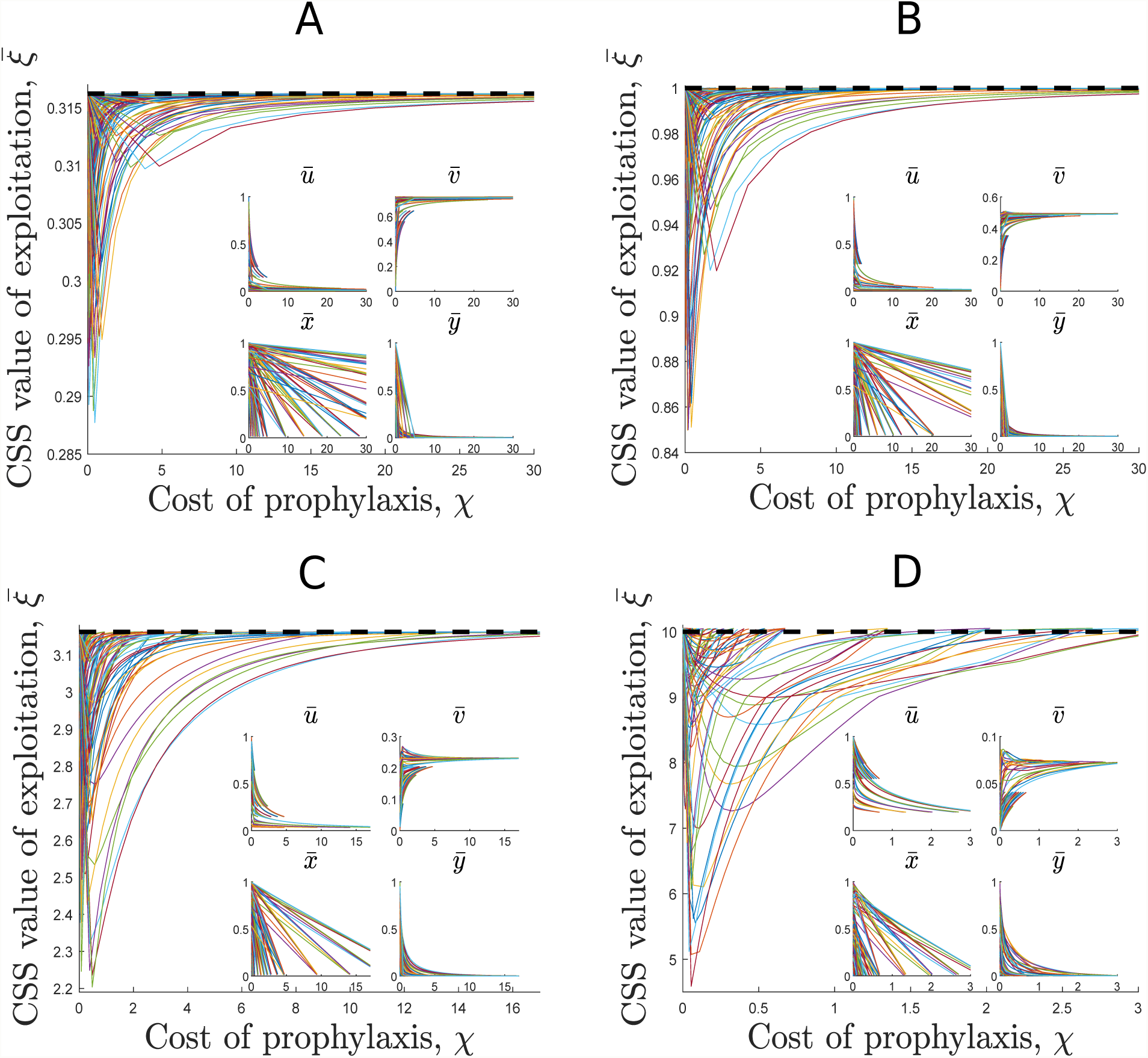
Plots of the CSS level of pathogen exploitation found using the numerical procedure described in the main text. Panel A contains results for *κ* = 0.1, panel B for *κ* = 1, panel C for *κ* = 10, and panel D for *κ* = 100. Inset figures show the equilibrium values of the epidemiological variables *u, v, x*, and *y*. The cost values presented are all below the critical cost threshold *χ*_*c*_. In all cases, the CSS level of exploitation is lower than the benchmark value 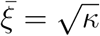, plotted as a dashed black line, observed in the absence of prophylaxis.

This pattern holds for a wide range of parameter values chosen to satisfy the conditions described in the previous section. Specifically, we investigate four different values of *κ*: *κ* = 0.1, *κ* = 1, *κ* = 10, and *κ* = 100. For each of these values, we choose five values of *β*_max_ above the 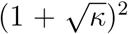 threshold: 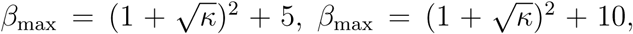, 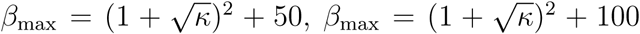 and 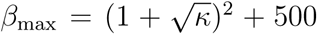. We then choose five values of *k* (*k* = 1*/*(𝓡_0_ − 1) + 5, *k* = 1*/*(𝓡_0_ − 1) + 10, *k* = 1*/*(𝓡_0_ − 1) + 50, *k* = 1*/*(𝓡_0_ − 1) + 100, and *k* = 1*/*(𝓡_0_ − 1) + 500) and five values of *ε* spread evenly between the threshold 𝓡_0_*/*((*k* +1)(𝓡_0_ −1)) and 1, for a total of 500 combinations of parameter values.

Although the decline in the CSS value of exploitation shown in figure 2 appears modest, recall that one time unit is equivalent to the average lifetime of an individual in the population. This means that the change in the duration of infection as 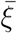 changes is on the order of years. For example, if the average lifespan of an individual in the population is 79 years, then a decrease from 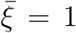 to 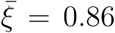 (as seen in figure 2, panel B) corresponds to an increase in the duration of infection of approximately three years.

## 4. Discussion

We study the impact of measures taken by hosts to limit disease transmission. Here, the willingness among hosts to engage in these prophylactic behaviours responds to changing costs and benefits. We focus on long-term evolution of a pathogen, defined by successive mutations until an equilibrium state is reached [22], alongside the rapid evolution of host behaviour. We find that, provided the cost of prophylaxis is below a critical value, hosts engage in the associated protective behaviour with positive frequency. In addition, when protective behaviour among hosts does occur, pathogen virulence — reflected by the evolutionarily stable exploitation level — is always lower than it is in the absence of prophylaxis. Finally, we find that stable virulence is lowest for intermediate frequency of protective behaviour among hosts (indirectly, intermediate cost of prophylaxis).

This study contributes to the growing body of work that shows host behaviour, in general, influences pathogen evolution. Much of this work has considered vaccination behaviour, in particular, and has described both beneficial and detrimental evolutionary outcomes. In the case of human papillomavirus (HPV), for example, theoretical work predicted HPV vaccination will select for higher levels of virulence [13]. By contrast, empirical evidence suggests that vaccination can actually limit the ecological opportunity open to certain HPV types [33]. In keeping with the mixed nature of results, Gandon et al. [34, 35] find that the direction of selection acting on pathogen virulence depends on the mechanism by which vaccination works.

More closely related to the current study are the conclusions of Pharaon and Bauch [16]. They show that host prophylactic behaviour in response to an endemic disease can allow the invasion of a pathogen strain that is more virulent than the resident. Our model predicts that such an outcome cannot occur as a result of the fact that our benchmark result, established in the absence of prophylactic behaviour, is our worst case scenario. Consider a mutant pathogen with a virulence 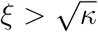. When the cost is above its critical value so that no one is engaging in prophylaxis, then this mutant cannot invade a resident population at the CSS 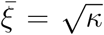. If we then decrease the cost below its critical value so that individuals begin to take prophylactic measures, the CSS exploitation level decreases away from 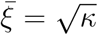 and so the mutant is still unable to invade the resident population.

The discrepancy between our prediction and those of Pharaon and Bauch is due to the differences in how we model the dynamics of host behaviour. In addition to looking at actual risk instead of perceived risk (as in [16]), we track both susceptible and infective protectors. Moreover, the evolution of host behaviour in our work is not governed solely by the replicator dynamics. Our model consists of additional terms to represent the effects of births and infection on the proportion of individuals engaging in prophylactic behaviour. These key differences are missing from previous work [16] and lead us to the conclusion that, for all feasible sets of parameter values, we expect a decrease in pathogen exploitation away from the benchmark value found in the absence of prophylactic behaviour.

To get some intuition into the decrease in virulence associated with protective host behaviour, we first recall the two tradeoffs faced by a pathogen in our model. Since the fitness function (9) is the product of the average duration of infection and the rate of production of new infections, changing virulence alters the balance between these two quantities. Changing virulence also alters the trade-off between creating new infections in protectors versus non-protectors.

When the cost of prophylaxis is high, the balance between creating new infections in different kinds of hosts depends strongly on virulence. Consequently, a mutant pathogen can effect a shift in the relative proportions of new infections it creates in protectors and non-protectors, respectively. By decreasing its virulence, a pathogen contributes to a reduction in the force of infection felt by the average individual in the population. This, in turn, provides hosts with an incentive to shy away from prophylactic behaviour. Essentially, a reduction in virulence gives hosts the excuse they need to stop engaging in the costly behaviour that interferes with disease transmission. In a way, our result is reminiscent of other pathogens that control host behaviour for their own gain. While our model pathogens are not directly controlling hosts like the pathogen *Ophiocordyceps unilateralis* does with the ant *Camponotus leonardi* [36] (or *Schistocephalus solidus* with the stickleback fish *Gasterosteus aculeatus* [37], or *Leucochloridium paradoxum* with the snail *Succinea putris* [38]), ours do indirectly influence the hosts’ economic agency.

As the cost of prophylaxis is reduced, the frequency with which mutant new infections are found in one kind of host or another becomes less dependent on virulence. Eventually, this frequency becomes totally decoupled from virulence when the cost is zero. For lower cost, then, the pathogen is only weakly able to influence the type of new infections it creates. Instead, it must fall back on influencing the trade-off between duration of infection and rate of production of new infections. This more classical trade-off has been studied in previous work, e.g., [8], and the standard prediction is that natural selection will push for increased virulence to decrease the duration of infection and maintain balance with a slowing transmission rate. As the cost of prophylaxis is decreased and the trade-off between duration of infection and rate of production of new infections becomes dominant, this standard result then explains the increasing virulence seen in our model. In this way, our results also contribute to the larger body of research looking at pathogen trade-offs by incorporating this nested trade-off where the pathogen is able to influence the type of infection created in addition to the more classical trade-off between transmission and recovery [27, 26].

As with any modelling endeavour, we have made some simplifying assumptions to reduce the mathematical complexity of our model. For example, we have assumed that individuals instantly update their behaviour when receiving new information about the progression of the disease. In reality, there is a time delay between receiving information and deciding to modify behaviour. Previous work has studied the effects of including a delay in the form of waning immunity either independently [39] or together with a delay in the form of a latent period following infection [40], and found that this can create periodicity in the model solutions. Other work has also investigated the effects of these delays on effective vaccination strategies [41]. Future work could extend our model to include the lag in information gain and behaviour modification, with the expectation that this would cause oscillations in the predicted virulence level.

One could also relax the assumption that individuals sample from all other individuals in the population when deciding whether or not to take protective measures. Epidemic models have previously been extended to include spatial structure through the use of networks, with different types of networks providing qualitatively different predictions [42]. Others have studied the interaction between network models and host heterogeneities in the case of sexually transmitted diseases [43], and the effects of adaptive networks where individuals may build and sever connections during the progression of a disease [44]. Networks could be incorporated here to explore the decoupling of social interactions related to disease transmission and those related to information transmission. Based on previous work [45], it is expected that this decoupling could lead to a lower level of pathogen virulence.

It is tempting to use our evolutionary predictions to inform public-health policy. While pathogen virulence does reach a minimum value for an intermediate level of cost of prophylaxis, our results (figure 2) also show that lowering the cost even farther below this level leads to fewer infections even if those infections are from a more virulent pathogen strain. Moreover, the pathogen’s virulence is always below the level predicted in the absence of individuals engaging in prophylactic behaviour. There is a balance, then, between the level of cost that is optimal for minimizing prevalence and the level optimal for minimizing pathogen virulence. It is important to recognize that efforts to minimize the cost of prophylaxis will result in a pathogen strain that is more virulent than it otherwise might be. More broadly, our work suggests that conversations about disease management and infection should be more inclusive towards the effects of human behaviour.

## Funding

This work was supported by an NSERC-CRSNG Discovery Grant (RGPIN: 2019-06626).

## Appendix A: Linear Stability Analysis

The full resident system of our model (5) in the main text is built on the standard twodimensional SIR model. This standard two-dimensional model has a Jacobian matrix of

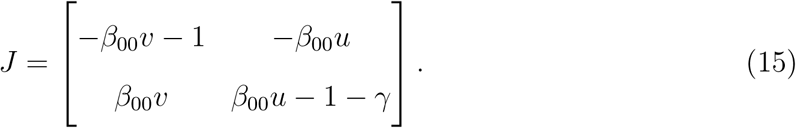

which, when evaluated at the disease-free equilibrium (DFE) 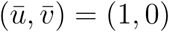, has eigenvalues *λ*_1_ = −1 and *λ*_2_ = *β*_00_ − 1 *-γ*. The DFE is stable when both eigenvalues are negative, which leads us to the condition that 𝓡_0_ = *β*_00_*/*(1 + *γ*) *<* 1 for stability.

When 𝓡_0_ exceeds this threshold, the DFE becomes unstable and the standard two-dimensional system moves towards an endemic equilibrium 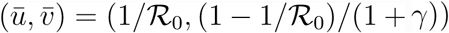. To determine the region in which this equilibrium remains stable after we incorporate the host behavioural dynamics, we need to consider the Jacobian matrix of our four-dimensional system (5) evaluated at 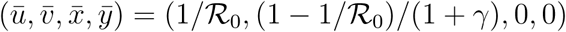:

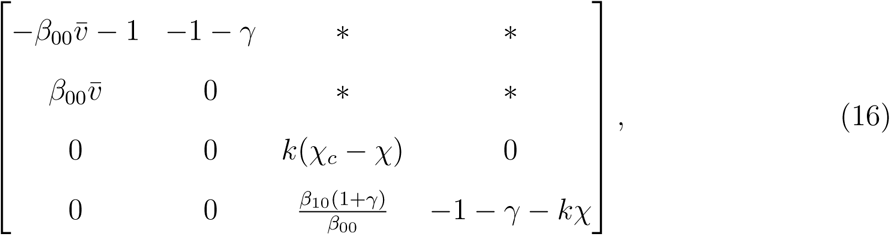

where 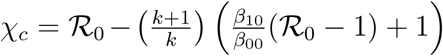 and asterisks denote entries that are possibly non-zero. Since this matrix is block upper triangular, the eigenvalues are given by the eigenvalues of the 2 *×* 2 matrices on the diagonal. The 2 *×* 2 matrix in the upper left is the Jacobian matrix of the standard two-dimensional SIR model evaluated at the endemic equilibrium 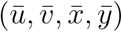, which we know has negative eigenvalues whenever 𝓡_0_ *>* 1. The 2 *×* 2 matrix in the bottom right is lower triangular, so its eigenvalues are the entries on the main diagonal. The second of these entries is always negative, while the first is negative as long as *χ > χ*_*c*_. This defines a critical cost threshold where for *χ > χ*_*c*_ the endemic equilibrium 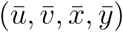 is stable, while for *χ < χ*_*c*_ our system tends towards an endemic equilibrium that contains protectors in some non-zero quantities.

## Appendix B: Perturbation Analysis

Letting **r** = (*u, v, x, y*) and *s* = *ξ*, we can express the four equations governing the epidemiological dynamics of our system as the vector-valued function **F**(**r**, *s*; *χ*) and the equation describing the evolutionary dynamics of pathogen exploitation as the scalar-valued function *G*(**r**, *s*; *χ*). We know that below the critical cost threshold *χ*_*c*_ the endemic equilibrium 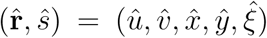 is stable, while above this threshold our system tends towards the protector-free equilibrium 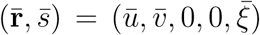. The critical cost level represents a bifurcation point where these two equilibria coincide and undergo an exchange of stability. To study how our system reacts as we decrease the cost away from this threshold, we introduce a perturbation parameter *δ* = *χ − χ*_*c*_ and take a first-order approximation to our new equilibrium point 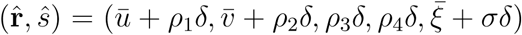. Knowing that this equilibrium point must satisfy 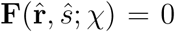 and 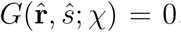, our goal is to find expressions for the perturbation coefficients *ρ*_1_, *ρ*_2_, *ρ*_3_, *ρ*_4_, and *σ*.

If we treat 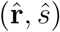 as a function of *χ*, we can make a first-order Taylor series approximation centred around *χ*_*c*_:

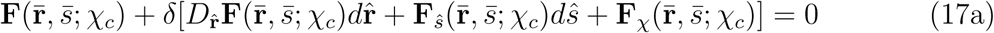

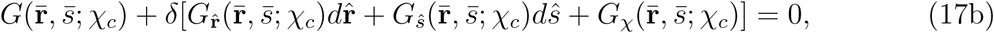

where subscripts denote partial derivatives and 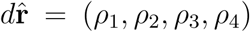 and *dŝ* = *σ* are the derivatives with respect to *χ* of 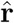 and *ŝ*, respectively. We know that 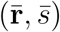 is an equilibrium point, so the first term in equations (17a) and (17b) evaluates to zero. We also observe that every term in **F** and *G* involving *χ* is multiplied by at least one of *x* or *y*, and so the partial derivatives with respect to *χ* vanish when we evaluate at 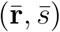. This simplifies (17) to:

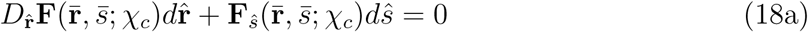

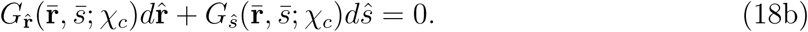

We can write (18) more succinctly as 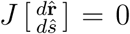 where the matrix *J* has the following structure:

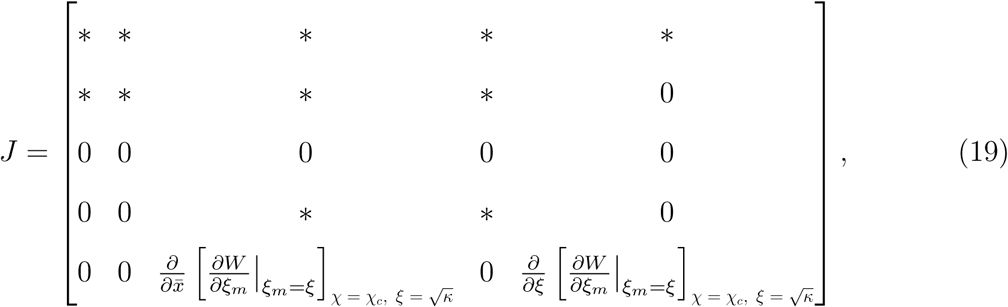

with asterisks denoting entries that are possibly non-zero. Since *J* is a block triangular matrix, the eigenvalues are given by the eigenvalues of the matrices on the main diagonal. The 2 *×* 2 matrix in the upper left is the Jacobian matrix arising from the linearization of the standard SIR model around the protector-free endemic equilibrium. Since the lower-right 3 *×* 3 block is lower triangular and has a zero entry on its main diagonal, we can see that zero is an eigenvalue of *J*. This allows us to interpret 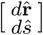 as the eigenvector of *J* associated with the zero eigenvalue, and so shows that there is a non-trivial solution for our perturbation coefficients.

While an analytic expression for this eigenvector can be found, it is unwieldy. Of more interest is the sign of the perturbation coefficient *σ*, as this tells us in which direction 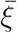 moves as we decrease the cost below its critical value. The third row of (19) tells us that 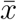 is a free variable, and the last row tells us that there is a simple relationship between this free variable and 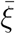. In particular, if we consider finding the eigenvector 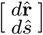 by solving the expression 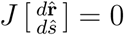, then the last row of (19) tells us that

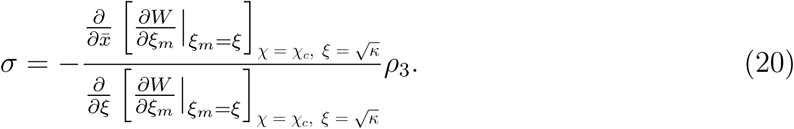

We know that the denominator of (20) is always negative since 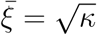 is convergence stable (see (11)), and we know that *ρ*_3_ *>* 0 since the proportion of susceptible protectors increases as the cost is decreased below its critical value. It follows, then, that the sign of *σ* is controlled only by the numerator of (20) and so we arrive at equation (14) in the main text.

## Appendix C: Maple Code

Here, we present the Maple code, described in section 3(b) of the main text, used to check the sign of the perturbation coefficient *σ* for a range of parameter values near the critical cost threshold. Using *δ* = *χ − χ*_*c*_ as our perturbation parameter, we first define the differential equation for *y*_*m*_ and solve for the equilibrium value:

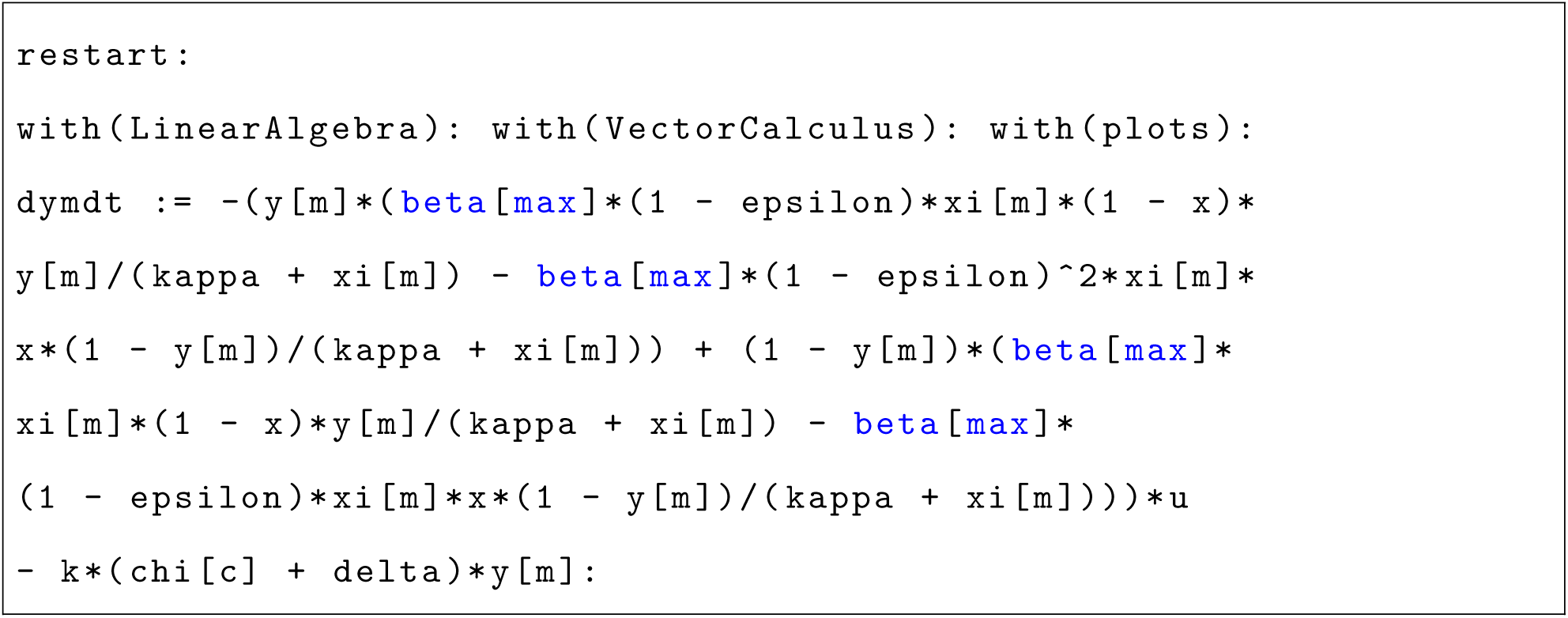

Solving this equation returns two possible equilibrium values, so we check the derivative of the differential equation to find the stable one:

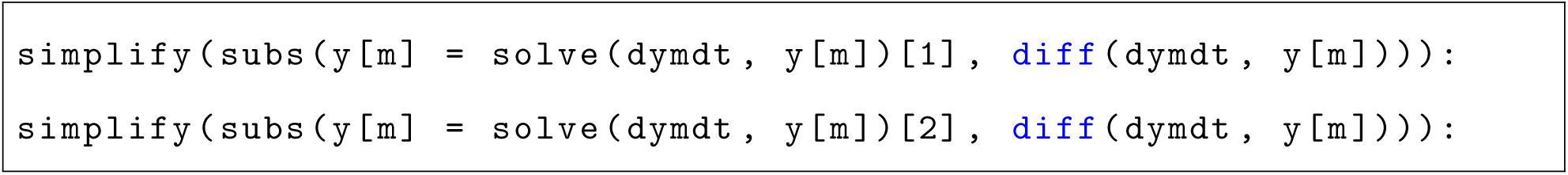

Since the first of these quantities is negative and the second is positive, this shows that the first root is the stable equilibrium:

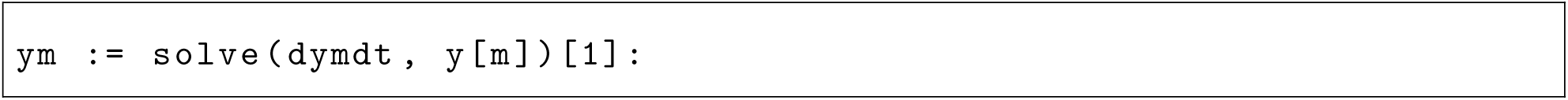

We now define the differential equations for *u, v, x*, and *y*, as well as the partial derivative of the payoff function with respect to the mutant pathogen exploitation level *ξ*_*m*_:

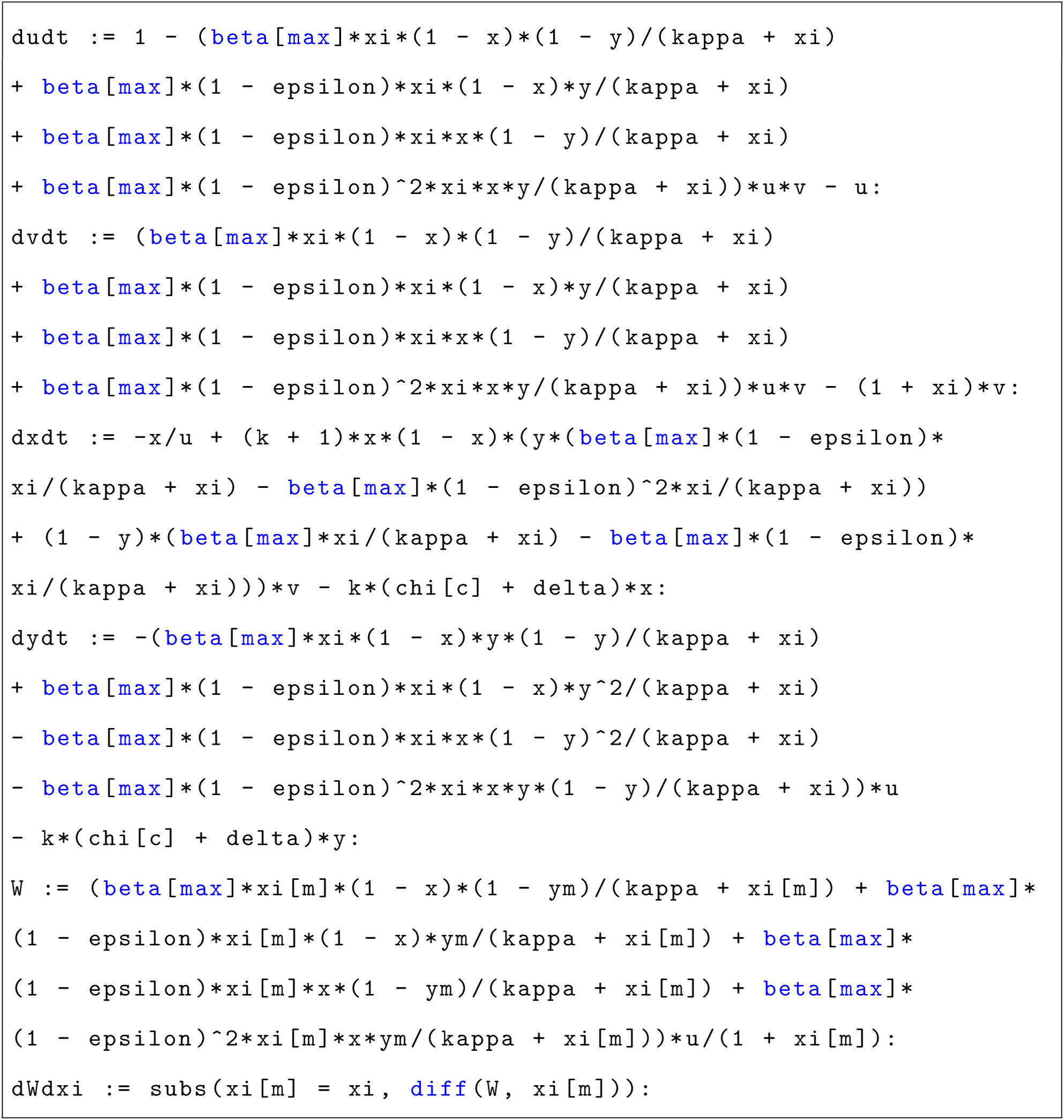

Using this, we put together the matrix *J* described in Appendix B:

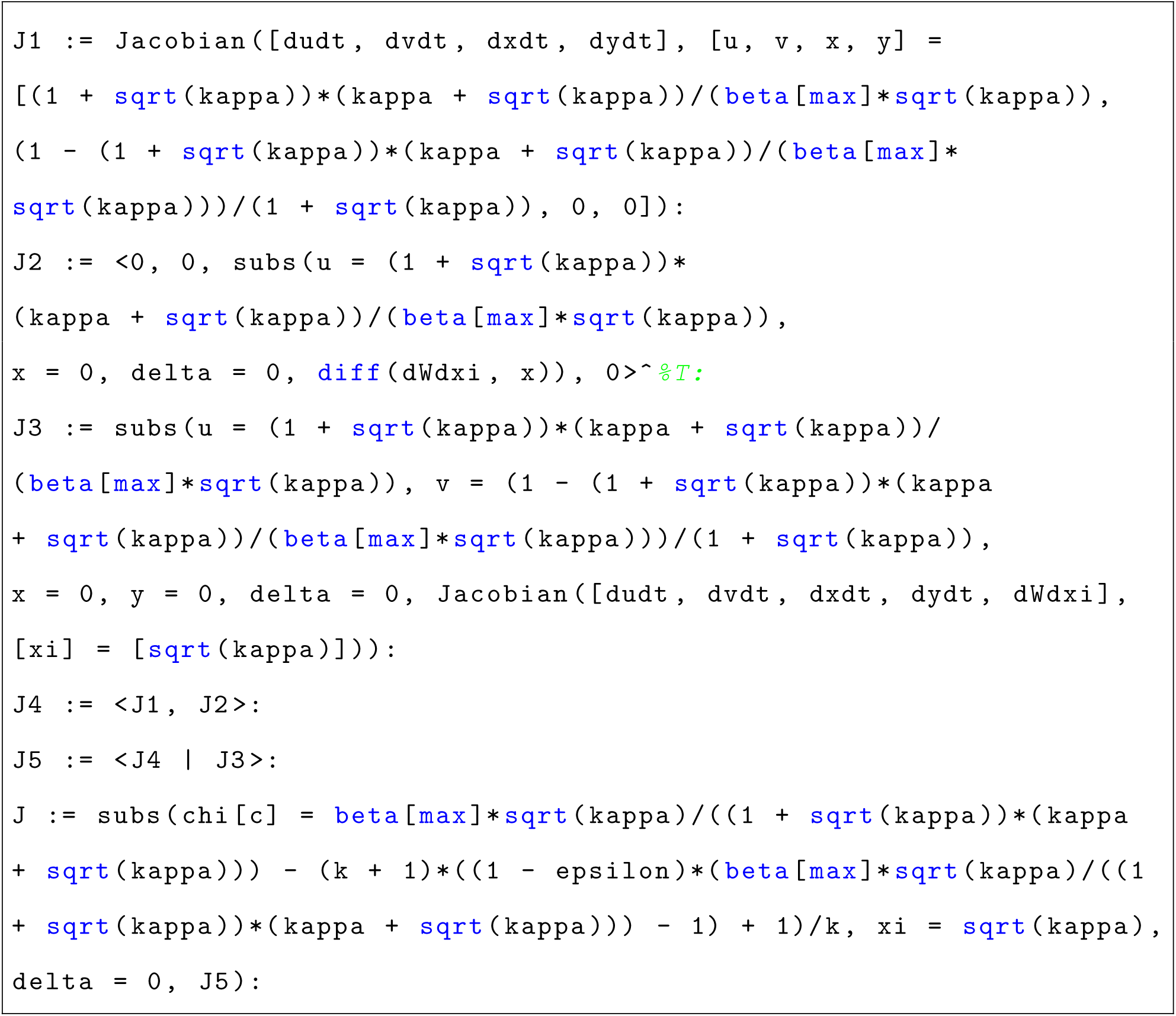

We then extract the entry of *J* corresponding to the numerator of *σ* in (14). We also define the critical cost threshold *χ*_*c*_:

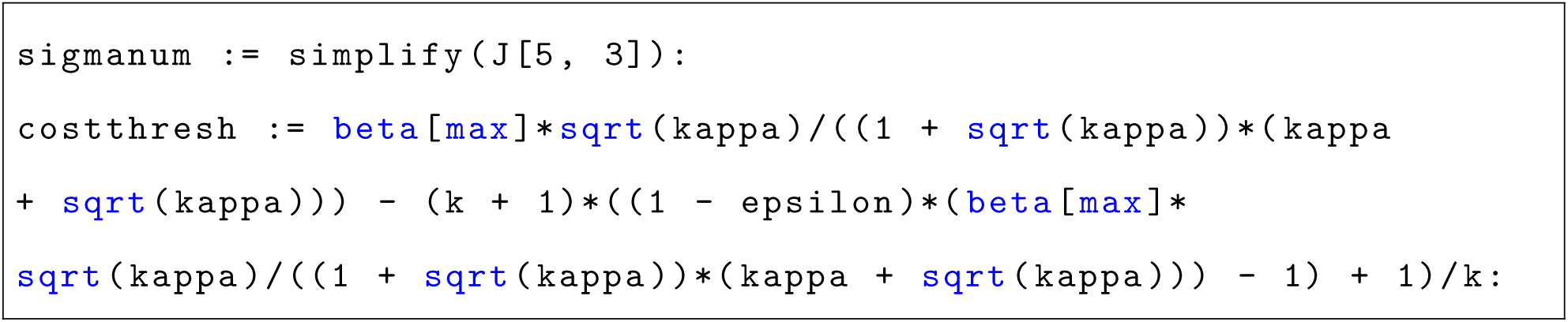

If we choose *β*_max_ and *κ* values so that 𝓡_0_ *>* 1, we get the following curve (where above the curve 𝓡_0_ *>* 1 and below the curve 𝓡_0_ *<* 1):

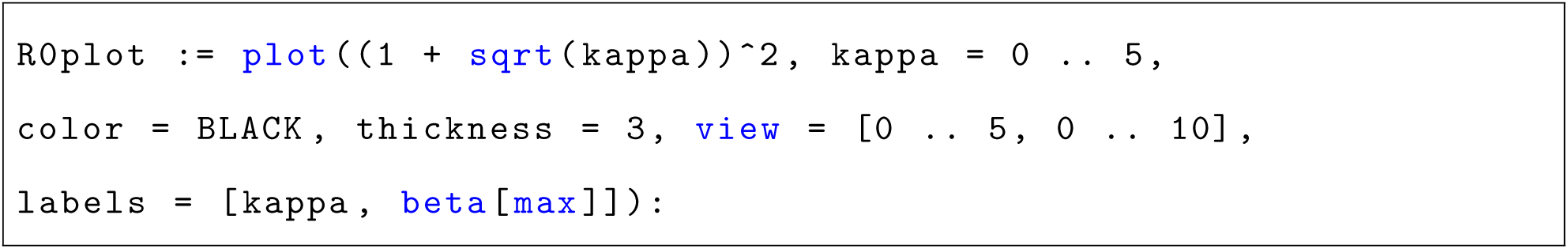

If we also choose *ε* values so that *χ*_*c*_ *>* 0 and *k* values so that *ε <* 1, we can generate a series of *β*_max_-*κ* curves and plot them together with the previous curve for 𝓡_0_:

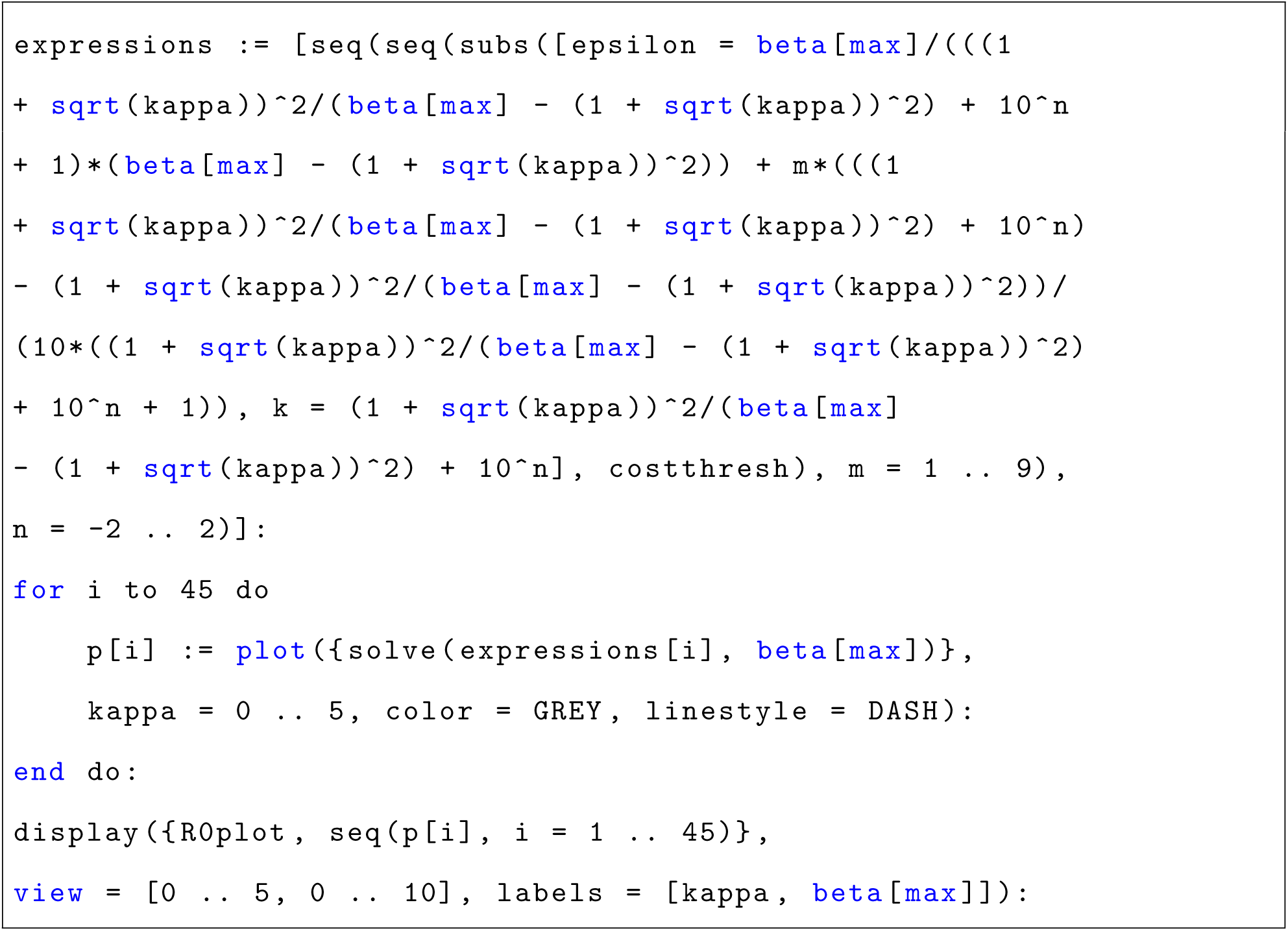

This generates the plot in panel A of figure 1 and suggests that all of the *χ*_*c*_ = 0 curves overlap with the 𝓡_0_ = 1 curve, meaning that the region in which 𝓡_0_ *>* 1 coincides with the region in which *χ*_*c*_ *>* 0. We can confirm this by looking at the difference between these two curves; running the following line of code will show that this difference is always exactly zero:

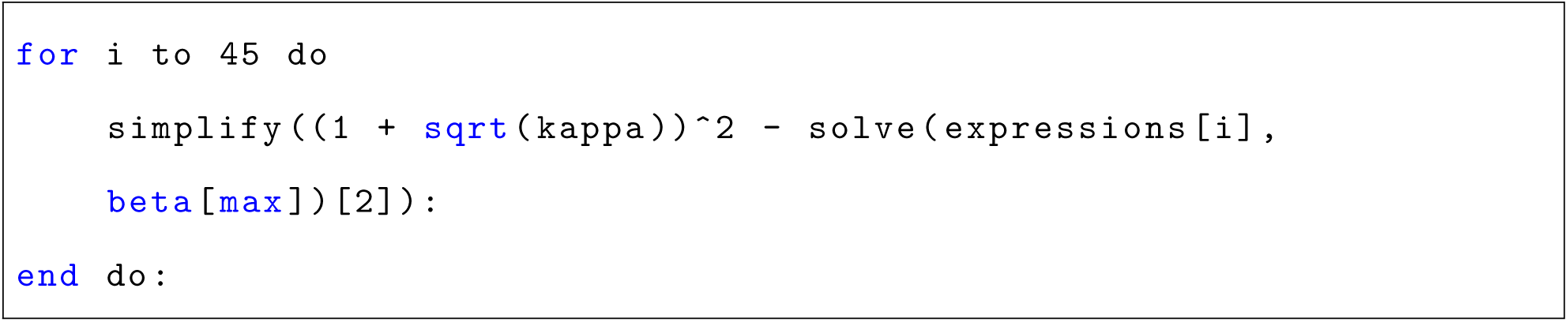

If we use the *ε* and *k* values chosen above, we can also look at the value of the numerator of *σ* as described in equation (14). This gives a second set of *β*_max_-*κ* expressions that we can plot over top of the expressions for *χ*_*c*_ and 𝓡_0_ above:

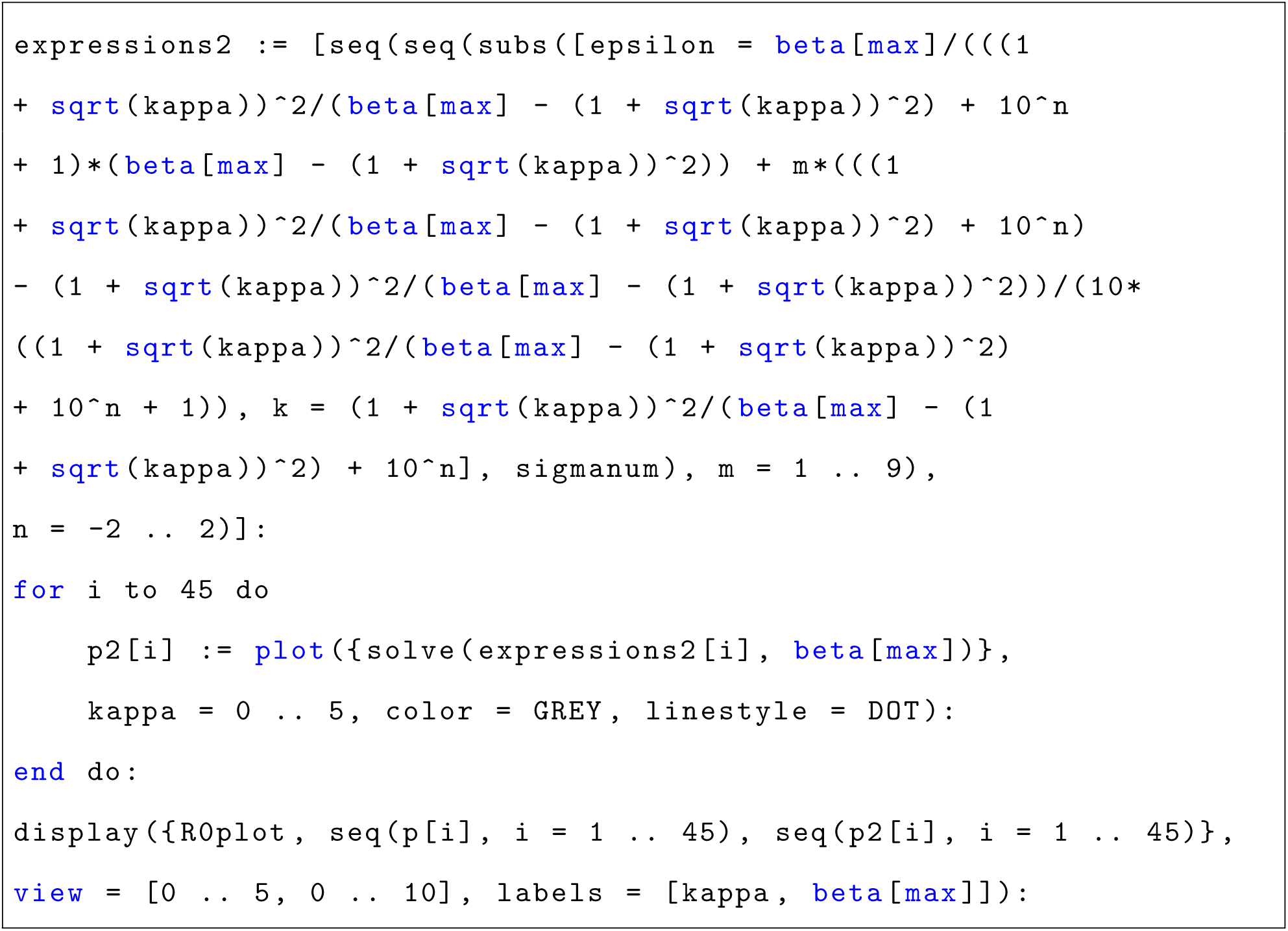

This produces the plot shown in panel B of figure 1 and shows that all of these curves that represent (14) equal to zero lay below the curve for 𝓡_0_ = 1. So for all sensible sets of parameter values (i.e., all parameter values that satisfy *χ*_*c*_ *>* 0 and 𝓡_0_ *>* 1), (14) has the same sign. We can take a test point in this region of parameter space to show that this sign is always negative, meaning that in a neighbourhood below the critical cost value the CSS pathogen exploitation level will decrease below its benchmark value.

## Appendix D: Matlab Code

Here, we present the Matlab code, described in section 3(c) of the main text, used to numerically find the CSS pathogen exploitation level. We start by defining functions for the transmission rates and the recovery rate:

Listing 1: Matlab function b00.m that computes the transmission rate *β*_00_(*ξ*).

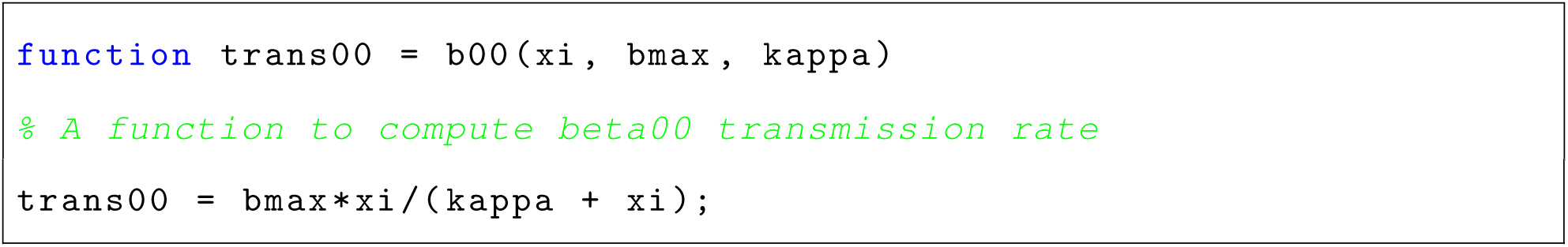

Listing 2: Matlab function b01.m that computes the transmission rate *β*_01_(*ξ*).

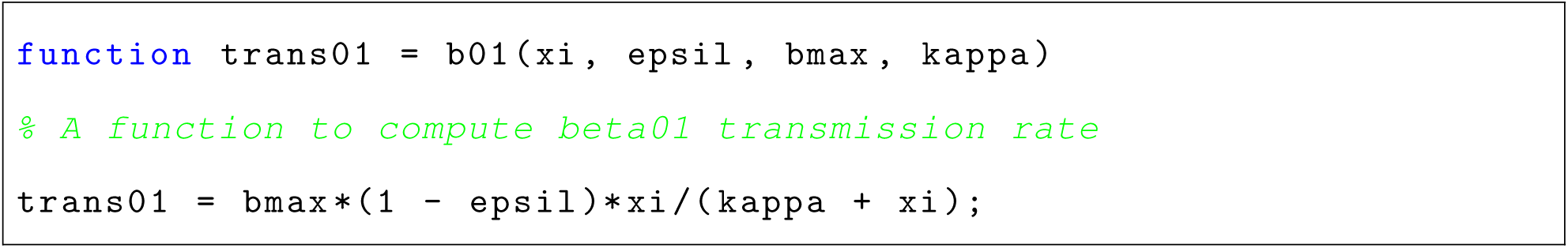

Listing 3: Matlab function b10.m that computes the transmission rate *β*_10_(*ξ*).

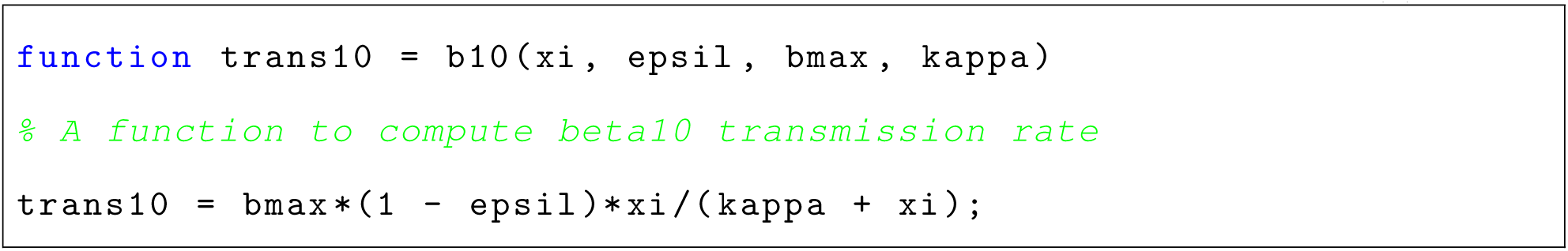

Listing 4: Matlab function b11.m that computes the transmission rate *β*_11_(*ξ*).

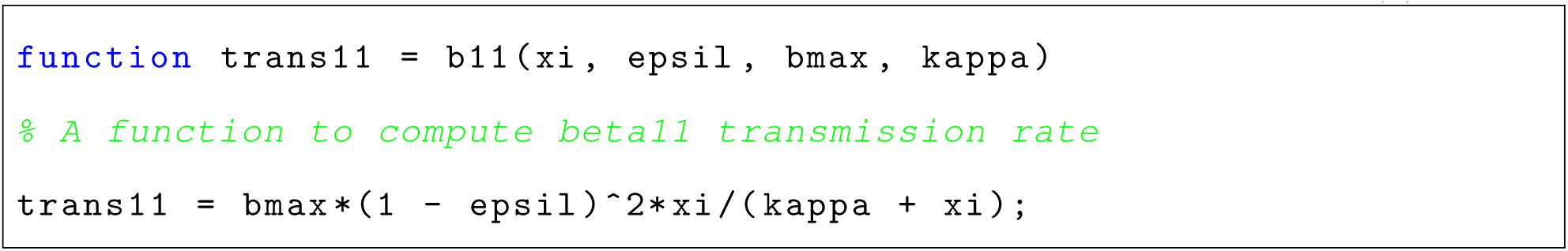

Listing 5: Matlab function g.m that computes the recovery rate *γ*(*ξ*).

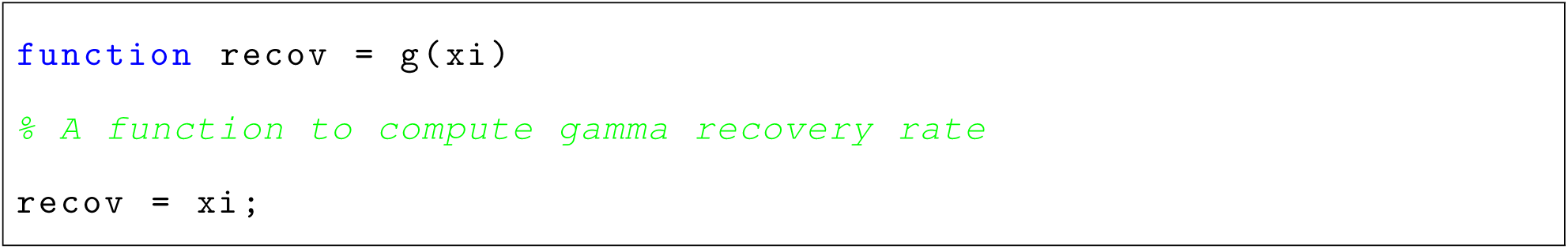

We also need functions to define the resident system and the partial derivative with respect to *ξ*_*m*_ of the fitness function:

Listing 6: Matlab function resident.m that computes the system of ODEs for the *u, v, x, y* system.

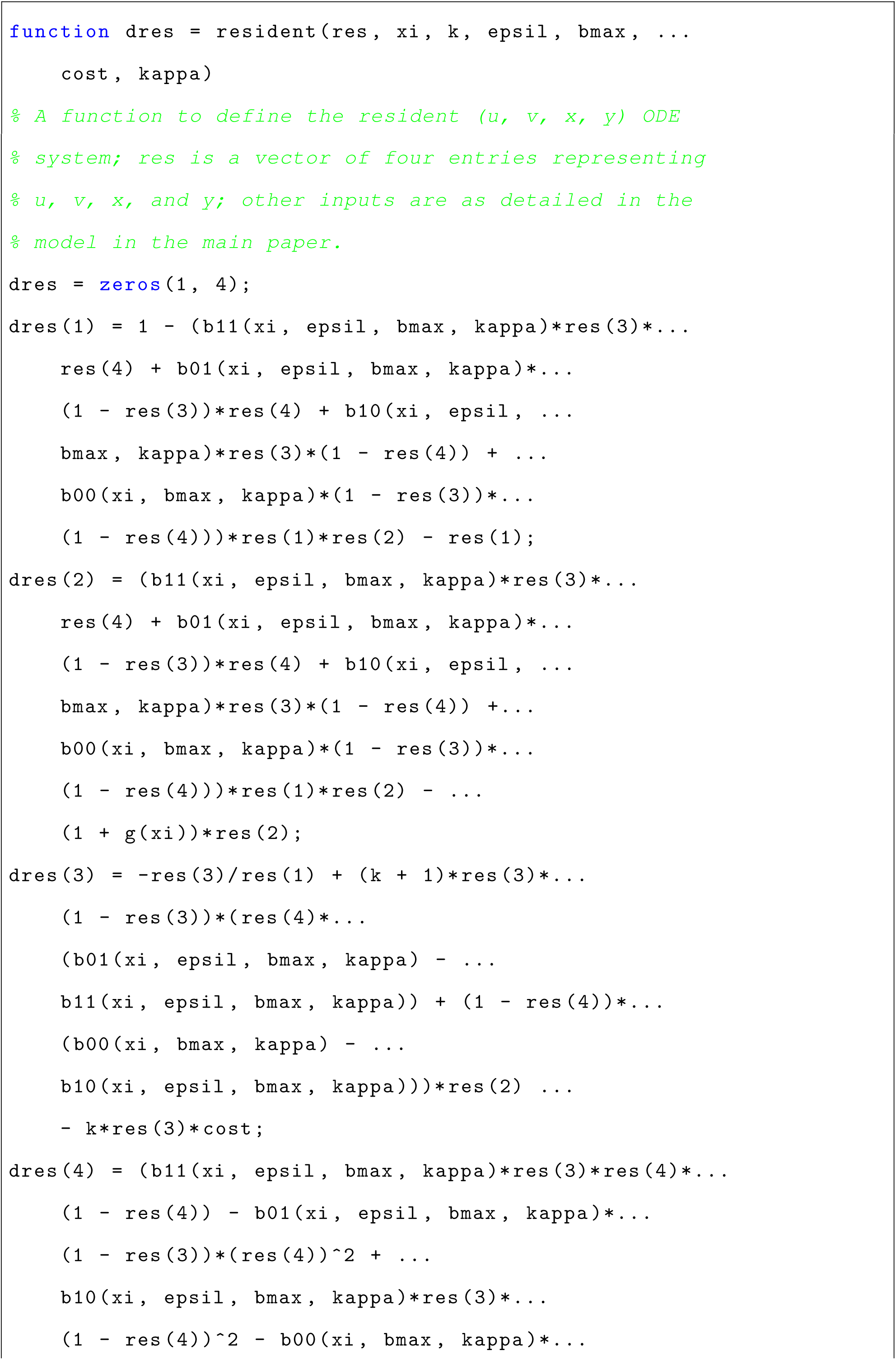

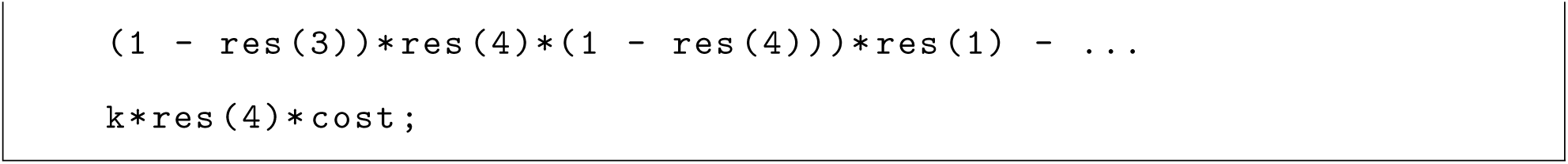

Listing 7: Matlab function dFitness.m that computes the partial derivative with respect to *ξ*_*m*_ of the fitness function.

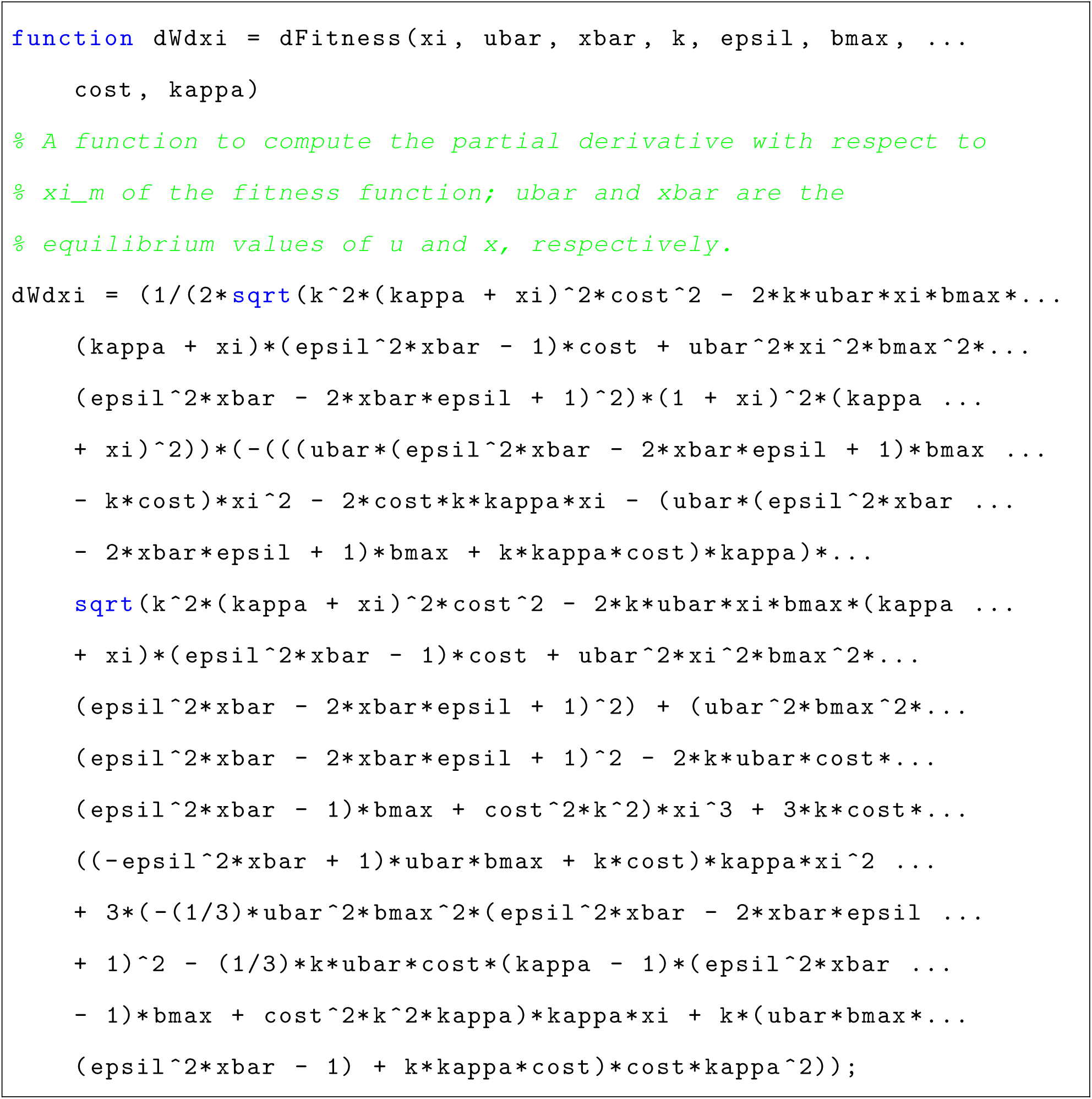

Finally, we use all of these to define a function that numerically approximates the CSS level of pathogen exploitation:

Listing 8: Matlab function findCSS.m that approximates the CSS level of pathogen exploitation.

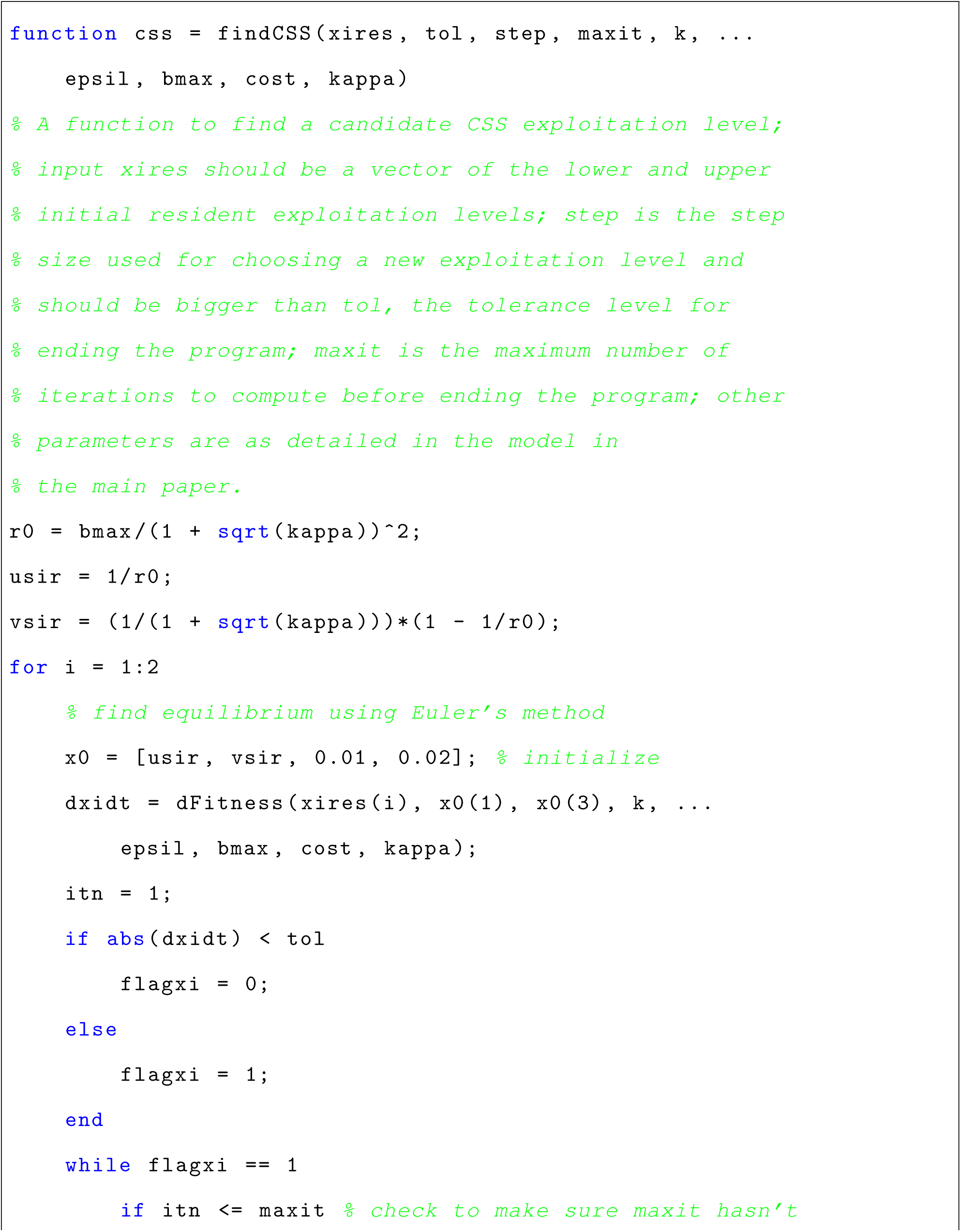

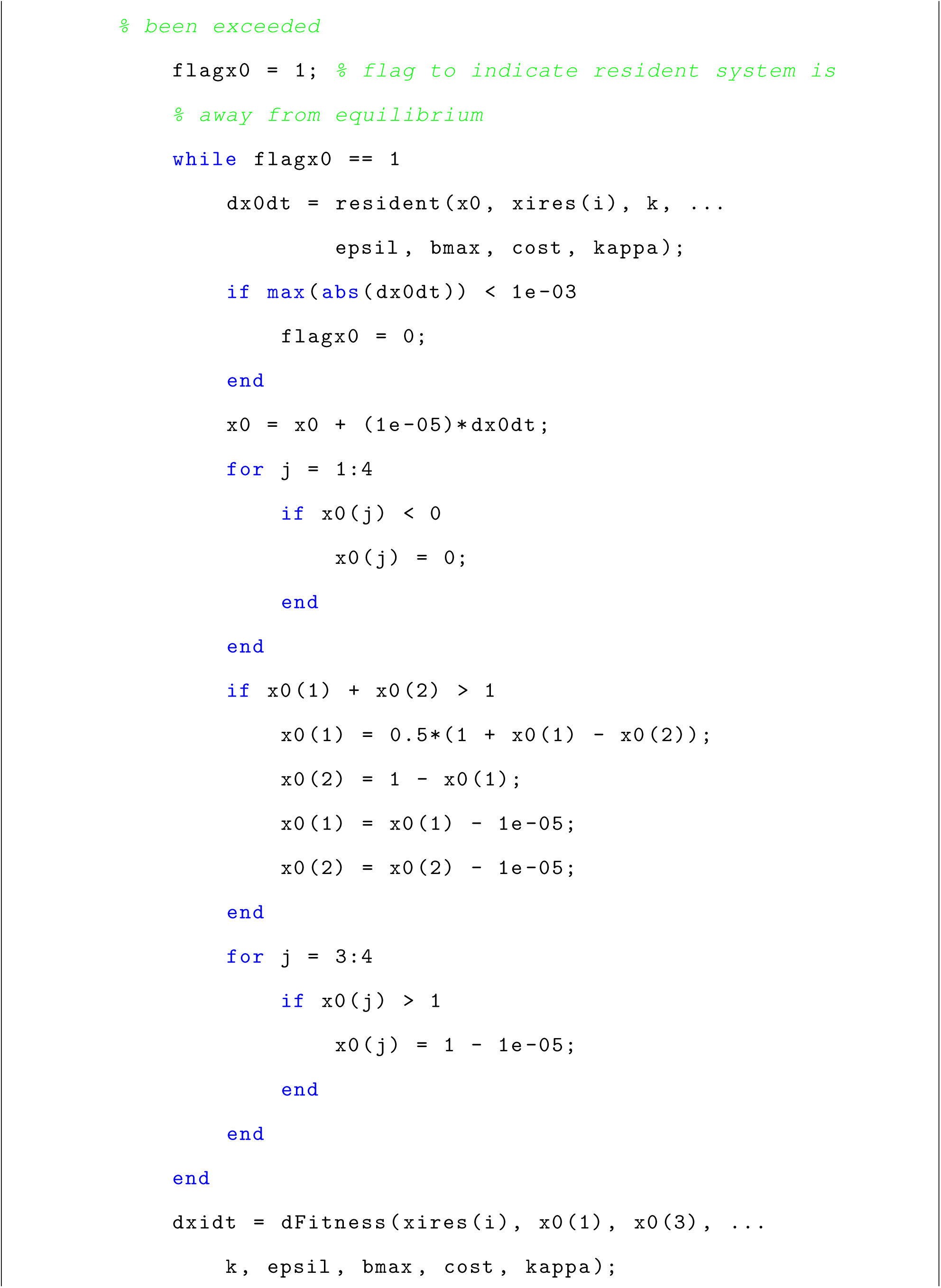

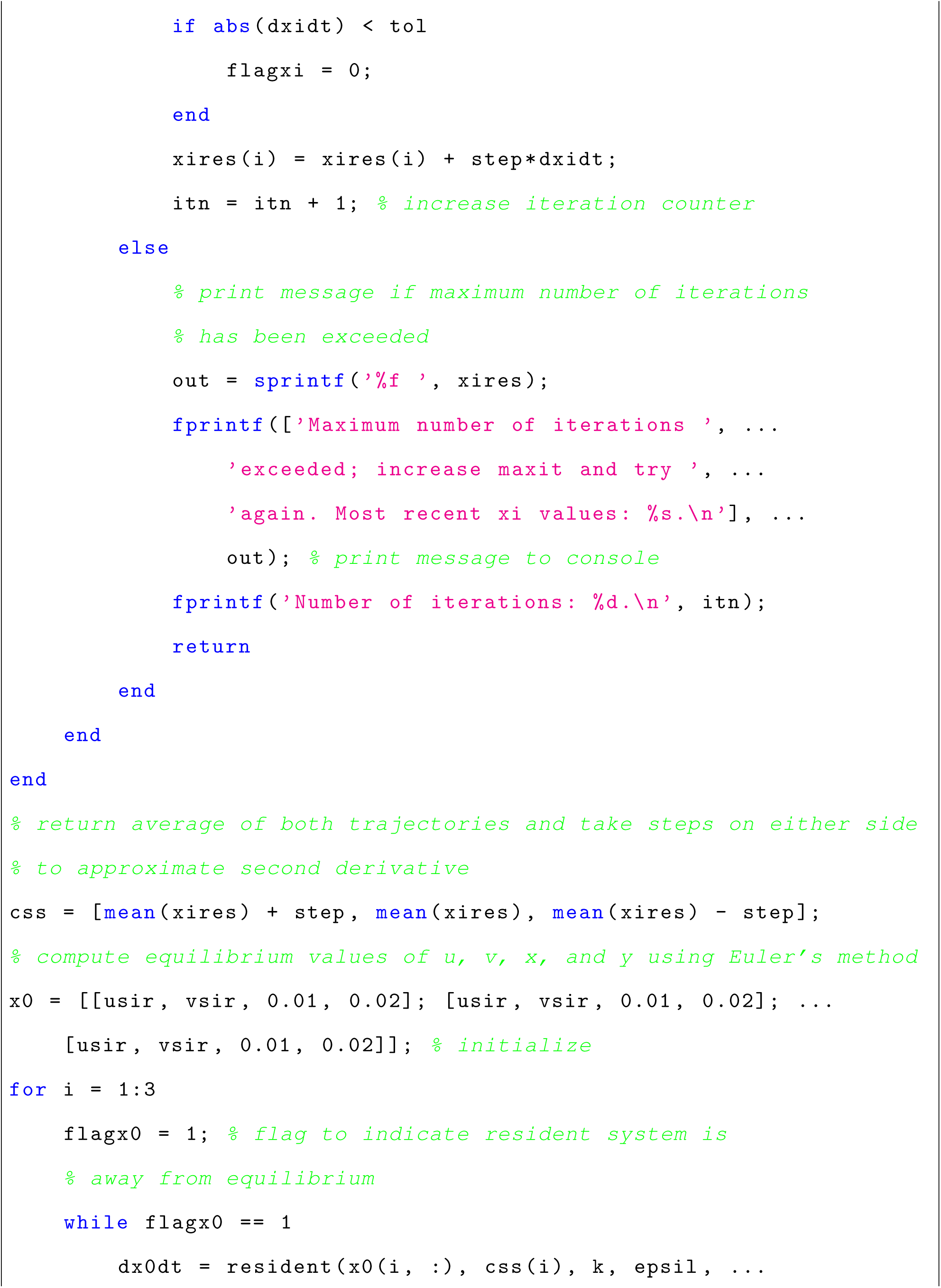

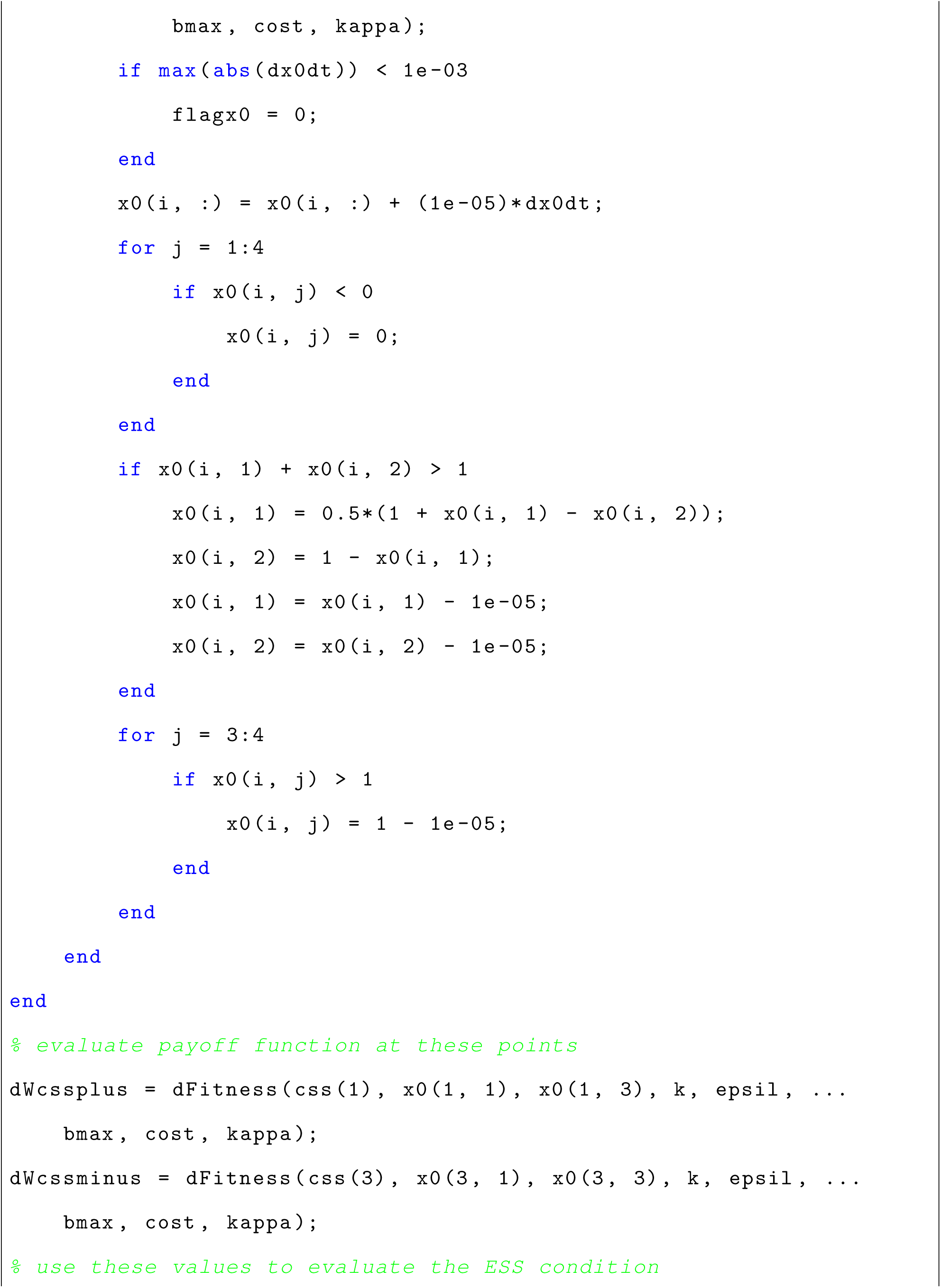

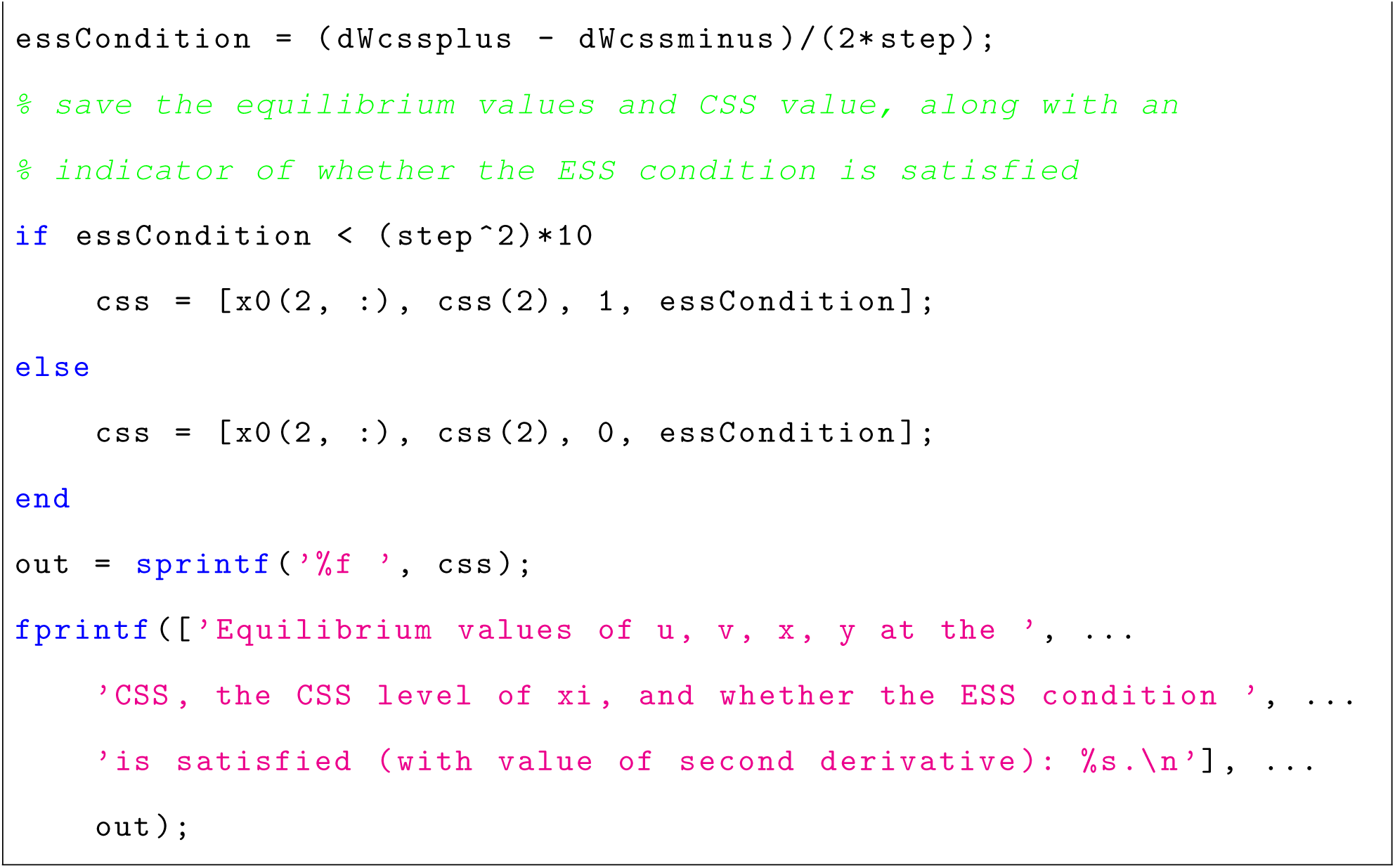

